# Lipid hydrogen isotope compositions primarily reflect growth water in the model archaeon *Sulfolobus acidocaldarius*

**DOI:** 10.1101/2024.10.24.620110

**Authors:** Carolynn M. Harris, Sebastian Kopf, Jeemin H. Rhim, Alec Cobban, Felix J. Elling, Xiahong Feng, Jamie McFarlin, Yuki Weber, Yujiao Zhang, Alice Zhou, Harpreet Batther, Ann Pearson, William D. Leavitt

**Author notes:** Address correspondence to Carolynn M. Harris. National Center for Science & Technology Evaluation, P.R. China.

## Abstract

The stable hydrogen isotope composition (δ^2^H) of lipid biomarkers can track environmental processes and remain stable over geologically relevant time scales, enabling studies of past climate, hydrology, and ecology. Most research has focused on lipids from the domain Eukarya (e.g., plant waxes, long-chain alkanes), and the potential of prokaryotic lipid biomarkers from the domain Archaea to offer unique insights into environments not captured by eukaryotic lipids remains unclear. Here, we investigate the H-isotope composition of biphytanes in *Sulfolobus acidocaldarius*, a model thermoacidophile and obligate heterotroph. We conducted a series of experiments that varied temperature, pH, shaking rate, electron acceptor availability, or electron donor flux. From these experiments, we quantified the lipid/water H-isotope fractionation (^2^ε_L/W_) values for core biphytane chains derived from tetraether lipids. The ^2^ε_L/W_ values are consistently negative (-230 to - 180 ‰) and are relatively invariant across all experiments despite a 20-fold change in doubling times and 2-fold change in lipid cyclization. The magnitude and relative invariance of ^2^ε_L/W_ values are consistent with studies on other heterotrophic archaea and suggests archaeal lipids may be faithful recorders of the δ^2^H composition of growth water. Our study highlights the potential of archaeal lipid δ^2^H as a hydrological proxy, offering new insights into environments where traditional proxies, such as plant-derived lipids, are not available, including extreme environments and extraterrestrial settings.

**Importance:** Reconstructing past climates is crucial for understanding Earth’s environmental history and its responses to changing conditions. This study examines *Sulfolobus acidocaldarius*, a thermoacidophilic archaeon that thrives in extreme environments like hot springs. These microorganisms incorporate hydrogen water in the growth environment into membrane lipids, creating hydrogen isotope signatures that can reflect hydroclimate conditions. Our findings show that these hydrogen isotope ratios remain consistent even under varying temperature, pH, oxygen levels, and electron donor fluxes, indicating a stable fractionation between lipids and water. This invariance suggests that *S. acidocaldarius* lipids could serve as a robust proxy for reconstructing ancient water H-isotope values, especially in extreme environments where traditional proxies, such as plant waxes, are absent. This research has broader implications for planetary-scale reconstructions, including potential applications in studying past climates on other planets, such as Mars, where similar microorganisms may have existed in hydrothermal conditions.

## 1. Introduction

The relative abundances of hydrogen isotopes (^2^H/^1^H) in water are set by physical, hydrological, and climatic factors (1, 2). Certain biomolecules incorporate these isotopes with a consistent offset (fractionation) relative to water, preserving a record of the isotopic composition (δ^2^H values) of the water present during growth (3). Some lipids can retain their original H-isotope composition for up to 10^8^ years and lipid hydrogen isotope compositions have been used to study hydrologic and environmental change over millennia to millions of years, dating as far back as the Paleoproterozoic (4–6). Most studies of biomarker H-isotope composition focus on lipids from photoautotrophic organisms (e.g., plant wax *n-*alkanes) or chemoautotrophic and heterotrophic bacteria (e.g., fatty acids) (3, 7–9) and the H-isotope compositions recorded in lipids from the Archaea are comparatively poorly understood. A small, but growing, body of literature, however, has begun to address this gap (10–15). Understanding the extent to which the δ^2^H value of archaeal lipids records the δ^2^H value of growth water requires a deeper examination of how environmental conditions impact this signal. A comprehensive investigation of the physicochemical factors affecting H-isotope fractionation between archaeal lipids and water (^2^ε_L/W_) is essential for employing archaeal lipid δ^2^H composition as a proxy for the δ^2^H composition of environmental water over space and time.

The ^2^ε_L/W_ value for a biomolecule integrates both the composition of source water and the net biosynthetic fractionations resulting from the specific pathways used during lipid biosynthesis. Reported ^2^ε_L/W_ values vary widely among domains of life. In phototrophic Eukaryotes (e.g., plants and algae), ^2^ε_L/W_ values of waxes (e.g., n-alkanes) are consistently negative, ranging from -150 to -50 ‰ (8, 16). It is important to understand the causes of this variability when applying lipid δ^2^H values to reconstruct water δ^2^H values. Net biosynthetic fractionation is generally consistent across taxa, though genetic factors introduce some variance (17–22). Environmental factors such as temperature, salinity, light availability, and CO_2_ concentration, also influence ^2^ε_L/W_ values, possibly via their impact on growth and biosynthesis rates (16, 23–28). These effects are well-characterized in plants, and to a lesser extent in algae, and can be accounted for when the δ^2^H values of plant waxes are used to reconstruct past hydroclimate (8, 29, 30).

In contrast to the coherent magnitude and direction of ^2^ε_L/W_ values observed from phototrophic eukaryotes, the ^2^ε_L/W_ of bacterial fatty acids varies widely in both direction and magnitude (-400 ‰ to +300 ‰) (31–38). Bacteria that use different central energy metabolisms (e.g., chemoautotrophy, photoautotrophy, or heterotrophy) exhibit distinct ranges of ^2^ε_L/W_ values, which has been attributed to differences in metabolic pathways and the substrate used for growth (e.g., simple sugars vs. tricarboxylic acid (TCA) cycle intermediates). A recent study suggests that variations in ^2^ε_L/W_ values from aerobic heterotrophic bacteria are explained by energy fluxes through the intracellular hydride carriers NADP⁺ and NADPH (38). Given this metabolism-dependent heterogeneity, the δ^2^H values of bacterial lipids in sediment records may have more utility as proxies for the energetic state of past microbial metabolisms (33, 36, 38).

Lipid-water ^2^H-fractionation from the domain Archaea have been understudied to-date. From what we do know, ^2^ε_L/W_ values have been reported for diether lipids (archaeol) in a mesophilic halophile (11) and a methanogen (13), tetraether-derived biphytanes in three thermoacidophiles and one thermomesophile (10, 15), and one aerobic marine chemoautotroph (14), and whole tetraethers or tetraether-derived biphytanes in several environmental sediment samples (10, 12). The observed ^2^ε_L/W_ values for these archaeal lipids are uniformly negative (i.e. -300 to -150 ‰) and similar in magnitude to those reported for isoprenoid lipids produced by eukaryotic phototrophs (e.g. C_15_ and C_30_ isoprenoids) (3, 25).

Our understanding of the factors controlling the H-isotope compositions of archaeal lipids are less developed than for the Eukarya and Bacteria. Recent experiments demonstrate that ^2^ε_L/W_ is nearly constant over a range of growth rates in a marine autotroph (*Nitrosopumilus maritimus* (14)) and that the impacts of carbon metabolism and growth phase on ^2^ε_L/W_ are minor (*Archaeoglobus fulgidus* (15)). This relative invariance in ^2^ε_L/W_ values is perhaps surprising considering the taxonomic and metabolic diversity of the investigated Archaea. The impact of environmental factors on this fractionation, however, has only been investigated within a single archaeal strain (*Haloarcula marismortui* (11)). Further investigation of the effects of multiple environmental factors on ^2^ε_L/W_ in Archaea is crucial, particularly since many archaea are adapted to extreme environments that can experience both rapid and large changes to physicochemical conditions (39).

Isoprenoid glycerol dibiphytanyl glycerol tetraethers (iGDGTs) are archaeal lipids with particular relevance to paleoenvironmental reconstructions (40). These tetraethers are structurally diagnostic, diagenetically robust, and routinely analyzed for paleoclimate reconstruction based on the relative abundances of specific structural moieties (40–42). Archaea can synthesize a unique series of iGDGTs that contain from zero to eight cyclopentane rings (e.g., iGDGT-0 through iGDGT-8), though the most highly ringed moieties (iGDGT-7 and -8) have only been recovered from hydrothermal spring environments (43). Members of the Thaumarchaeota phyla can also produce crenarchaeol, an iGDGT containing four cyclopentane rings and one cyclohexyl ring (43). The distribution of these moieties in natural systems is related to growth temperature and other environmental factors (44–48) and forms the basis of the TEX_86_ paleo sea surface temperature proxy (41). At low pH and high temperatures, Archaea synthesize more highly-ringed moieties as an adaptation to maintain membrane rigidity and reduce permeability. In hydrothermal systems, the ring distributions of thermoacidophilic archaea record the interactions between temperature, pH, energy availability, and other factors that influence growth rate (48–50).

To better understand the signals recorded by archaeal lipid δ^2^H composition in natural systems, it is necessary to determine the extent to which environmental conditions influence ^2^ε_L/W_ values. In this study, we determined the H-isotope fractionation between iGDGT-derived biphytanes and growth water (^2^ε_L/W_) for the thermoacidophilic and heterotrophic archaeon, *Sulfolobus acidocaldarius* (DSM 639). *S. acidocaldarius* evolved to grow optimally under multiple extreme conditions, namely low pH’s, high temperatures, and low oxygen partial pressure (51, 52). For this work *S. acidocaldarius* was selected as a model organism because it thrives in extreme environments devoid of traditional eukaryotic lipid proxies (e.g., plant waxes), and because it can grow under a wide range of physicochemical conditions, with rapid doubling times and high biomass yields (53).

Here we cultivated *S. acidocaldarius* over a range of physicochemical conditions that included shifts in temperature, pH, shaking rate, electron acceptor or electron donor flux. Changes in these environmental conditions are known to impact the growth rate and degree of iGDGT cyclization (47–49, 54, 55), but their impact on lipid/water fractionation is unexplored. Understanding how these environmental factors influence lipid/water fractionation is critical for interpreting archaeal lipid δ²H signatures in natural systems and refining their use as robust proxies for reconstructing past hydroclimate conditions.

## 2. Methods

### 2.1 Culture strains and growth

Axenic cultures of *S. acidocaldarius* (DSM 639) were grown aerobically under different environmental conditions (**Table 1**). Culture media was prepared following the Brock recipe (56) with the following modifications (see *Substrate* column in Table 1). For all experiments, media contained 5.9 mmol L^-1^ (0.2% w/v) sucrose as the organic substrate. The electron donor flux experiments were also supplemented with 4.0 mmol L^-1^ (0.1% w/v) NZ-amine in addition to sucrose.

**Table 1.** Summary of culture conditions and descriptive growth statistics for S. acidocaldarius grown over varying environmental conditions. δ^2^H_W_ is the H-isotope composition of media water at the time of lipid sampling. *Growth d*ata for batch and fed-batch environmental condition experiments were initially reported in Cobban et al., (2020), *growth* data for chemostat experiment was initially reported in Zhou et al., (2019). *For batch experiments, RPM is shaking rate; for bioreactor experiments, RPM is impellor speed.

Growth was monitored photometrically at regular intervals throughout each experiment by measuring the optical density (i.e., absorbance at 1 cm pathlength) of a culture aliquot at 600 nm (OD_600_) on a spectrophotometer. All experiments were inoculated to an initial OD_600_ between 0.004 and 0.008, using cells derived from an active pre-culture (grown on the same medium) in mid-exponential phase. Specific growth rates (division day^-1^; μ) were calculated using OD_600_ measurements during the early to mid exponential phase after Cobban et al., (2020). Doubling time (T_D_) was calculated as:

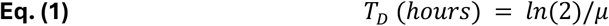

### 2.2 Experimental conditions experiments

We examined how temperature, pH, shaking rate, O_2_ availability, and electron donor (sucrose) flux influence H-isotope fractionation in *S. acidocaldarius* through a series of batch, fed-batch, and chemostat experiments.

#### Batch experiments

Three sub-experiments were performed in which one environmental condition (temperature, pH, or shaking rate) was systematically varied while holding the other two conditions constant (previously described in Cobban et al., 2020). The reference conditions were 70 °C, pH 3, and 200 RPM shaking rate. Shaking rate is considered a proxy for aeration rate, where faster speeds increase oxygen availability in the media, though it also increases physical agitation.

Batch experiments were performed in temperature-controlled, shaking incubators (Innova-42, Eppendorf) in 250 mL Erlenmeyer flasks containing 125 mL media covered with loose plastic caps to reduce evaporation. Evaporation was further reduced by maintaining a 2L tray of water in the incubator to humidify the atmosphere. All experiments were performed in quintuplicate. Biomass samples were collected in mid-exponential phase by centrifuging (3214 x *g*; Eppendorf 5810 R, S-4-104 rotor) 50 mL of culture for 15 minutes at 4 °C, the resulting cell pellets were stored at -80 °C until lipid extraction.

#### Fed-batch bioreactor experiments

To more precisely determine the impact of oxygen availability, we performed gas-fed batch experiments in bioreactors (previously described in Cobban et al., 2020). Briefly, *S. acidocaldarius* was grown in 300 mL of medium in 1 L glass vessels. To modulate availability (as flux) of the terminal electron acceptor O_2_ was introduced to bioreactors at four different partial pressures: 0.2, 0.5, 2.0 or 20 % (balance N_2_). The O_2_ dry mixing ratios 0.2, 0.5, and 2% were generated by mixing pure N_2_ mixed with 2% O_2_ at 9:1, 3:1, and 0:1 ratios, respectively, which was controlled via my-Control PID controllers (Applikon, Delft, Netherlands). The 20% experiment used a separate gas feedstock of Ultra ZeroGrade Air (AirGas). Culture conditions were maintained at 70 °C, pH 3, stirred at 200 RPM (Rushton-type impeller), and at a total gas flux of 200 mL min^-1^. Each experiment was performed in triplicate via three bioreactors operated in parallel. Biomass samples were collected by removing culture directly from the vessel, centrifuged at 4 °C, to pellet biomass, and then stored at -80 °C until lipid extraction.

#### Chemostat experiments

To determine the impact of electron donor flux, we performed chemostat experiments in which the dilution rate of the growth vessel was set to yield cell doubling times (T_D_) of 7.0, 21.3, and 40.0 hours (previously described in Zhou et al. 2020). Briefly, *S. acidocaldarius* was grown in 300 mL of medium maintained at 70 °C, pH 2.25, stirred at 200 RPM (Rushton-type impeller), and aerated with a constant flux of 200 mL min^-1^ Zero Air (ultra-high purity, UHP, AirGas/AirLiquide, Inc.). Three chemostats were operated in parallel under continuous culture conditions. When steady state was achieved (assessed via regular OD_600_ measurements), biomass samples were harvested by continually collecting the outflow of each bioreactor into a chilled vessel (0 to 4 °C) for between 1 and 4 days, followed by centrifugation at 4 °C and storage at -80 °C prior to lipid processing. Multiple extracts representing 2 to 3 sampling events at identical growth rates were combined before lipid extraction.

### 2.3 Lipid extraction and derivatization

Core lipid fractions (iGDGTs) were extracted from lyophilized cell pellets via acidic hydrolysis-methanolysis (45, 48). iGDGTs were chemically derivatized to biphytanes to permit the determination of the δ^2^H value of component biphytanes (10, 14). In brief, ether bonds were cleaved via digestion in 57% hydroiodic acid (HI) at 125 °C for 4 hours. The resulting alkyl iodides were hydrogenated (e.g., reduced to biphytanes, BPs) in H_2_/PtO_2_. An isotope dilution occurs when H is added during hydrogenation, which is corrected for during data processing and reduction (see Section 2.5).

### 2.4 Lipid identities & distributions

Isoprenoid BP chains were analyzed via GC-MS (Thermo ISQ LT with TRACE 1310) for compound identification and via GC-flame ionization detector (GC-FID; Thermo TRACE 1310) for compound quantification at the University of Colorado at Boulder Earth Systems Stable Isotope Lab (CUBES-SIL, Boulder, CO) (**FIG 1**). Biphytane yields and recovery were calculated using GC peak areas and co-injected n-alkane mixture as a quantification standard.

**Fig 1.**
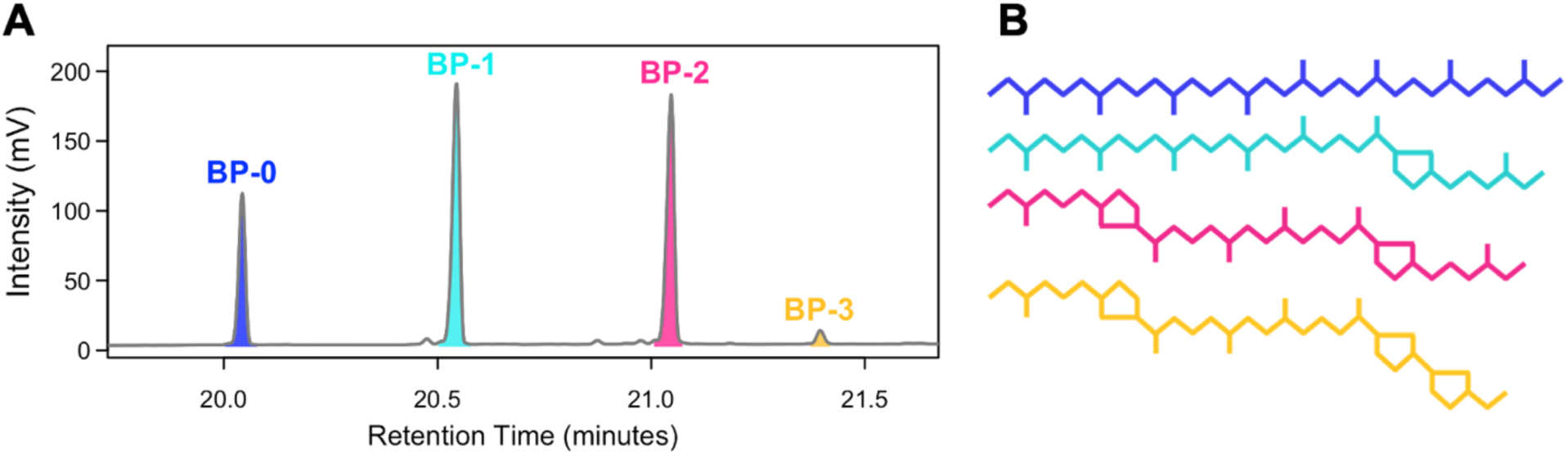
**(A)** Representative chromatogram of intensity vs. retention time for four iGDGT-derived biphytane (BP) moieties identified in *S. acidocaldarius* biomass that are named for the number of cyclopentane rings they contain. Although a BP-3 peak was detected, in many cases it is present in too low quantities to perform isotopic measurements **(B)** The identity and structure of BP-0 to BP-3.

To describe the average amount of cyclopentyl rings in each BP distribution, we calculated a BP-Ring Index based on the relative abundance of each BP moiety (**Eq. 2**). This metric is similar to the iGDGT-based Ring Index (**Eq. 3**), which describes the average amount of cyclopentyl rings in a distribution based on the relative abundance of each iGDGT moiety (48, 57).

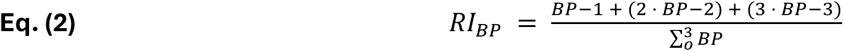

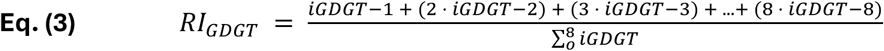

### 2.5 δ^2^H analysis of media waters, substrates, and biphytane lipids

#### 2.5.1 Media waters

Water samples for hydrogen isotope (δ^2^H) analysis were collected concurrently with harvesting biomass for lipid analyses. Aliquots of filter-sterilized growth medium water was measured for δ^2^H at the Stable Isotope Laboratory at Dartmouth College via a dual inlet isotope ratio mass spectrometer (IRMS; Thermo Delta Plus 304 XL) coupled to an H-Device (pyrolyzed to H_2_ gas) for water reduction by chromium powder at 850°C (previously described in Kopec et al. (2018). The δ^2^H_Water_ values were corrected using three laboratory standards spanning -161 to -7 ‰ vs. VSMOW, which were run every 8-10 samples. Analytical precision was <0.5 ‰.

#### 2.5.2 Substrates

The δ^2^H composition of non-exchangeable C-bound H from substrates used in growth media was analyzed in triplicate as trifluoroacetate derivatives via a Thermo Delta Plus XP coupled to a TC-EA (58). The δ^2^H composition of sucrose was -93.0 ± 1.2 ‰.

#### 2.5.3 Biphytanes

For all experiments, we report ^2^ε_L/W_ values for biphytane (BP) hydrocarbons derived via ether cleavage from acid extractable iGDGTs. The δ^2^H composition of individual BPs was determined by gas chromatography pyrolysis isotope ratio mass spectrometry (GC-P-IRMS) on a GC IsoLink II IRMS System (Thermo Scientific) at the University of Colorado at Boulder Earth Systems Stable Isotope Lab (CUBES-SIL, Boulder, CO). The system comprised a Trace 1310 GC fitted with a programmable temperature vaporization (PTV) injector and either a 30 m ZB5HT column (i.d. = 0.25 mm, 0.25 μm, Phenomenex, Torrance, CA, USA) or a 60 m DB1 column (i.d. = 0.25 mm, 0.25 μm, Agilent, Santa Clara, CA, USA), ConFlo IV interface, and 253 Plus mass spectrometer (Thermo Scientific).

#### 2.5.4 Data reduction and notation

The δ^2^H values of individual biphytanes (δ^2^H_BP_) were measured relative to H_2_ reference gas (δ^2^H_raw_) and calibrated to the international references scale (VSMOW) using a standard n-alkane mixture (A7, containing C15 through C30 n-alkanes spanning -9 to -263 ‰ vs. VSMOW; A. Schimmelmann, Indiana University). The A7 standard was combined with a C_36_ n-alkane (nC_36_, -259.2 ‰ vs. VSMOW; A. Schimmelmann, Indiana University) and measured throughout each analytical run at regular intervals and at different concentrations. The BP hydrogen isotope calibration was performed as in Leavitt et al., (2023) using the R packages *isoreader* (v 1.3.0) and *isoprocessor* (v 0.6.11) available at github.com/isoverse (59). In brief, δ^2^H_raw_ values were corrected for offset, scale compression and peak-size effects using an inverted multivariate linear regression which was applied to all standards and samples to determine δ^2^H_cal_ values. Further details of calibration and data reduction are available in Leavitt et al., 2023. The total analytical uncertainty of the corrected δ^2^H_BP_ values was calculated using standard error propagation of the peak-size adjusted error estimates and hydrogenation correction assuming all errors to be uncorrelated. The hydrogenation correction ranged from 10.3 to 12.7 ‰ and increased analytical uncertainty by up to 1.9 ‰.

All ^2^H/^1^H ratios are reported in delta notation (δ^2^H) in permil (‰) units relative to the international seawater standard on the VSMOW-SLAP (Vienna Standard Mean Ocean Water, Standard Light Antarctic Precipitation) scale.

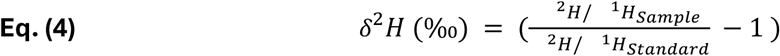

All fractionation factors are reported in epsilon notation (^2^ε) in permil (‰) units. The hydrogen isotope fractionation between growth water and BP lipids (^2^ε_L/W_) is calculated as:

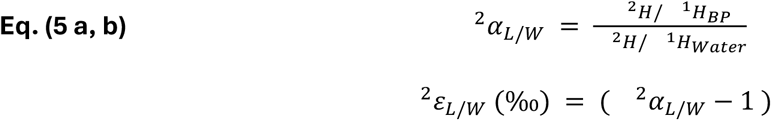

Corrected δ^2^H_BP_ values and resulting ^2^ε_L/W_ fractionation factors from biological replicates (n ≥ 1) and analytical replicates (n ≥ 3) were averaged for each experimental condition. All averages are weighted means of individual measurements (1/σ^2^ weights) to account for the amplitude-dependent range in uncertainties. The reported error estimate of each average is the larger of the standard deviation of all replicates or the propagated uncertainty from individual measurements. Abundance weighted averages for δ^2^H_BP_ and ^2^ε_L/W_ values include all BPs with relative abundance > 5%.

The ring difference (Δε/ring), or changes in ^2^ε_L/W_ with increasing number of ring structures is calculated for all pairwise combinations of ^2^ε_L/W_ values after Leavitt et al., (2023) where X and Y refer to the ring number of BP moieties and X > Y:

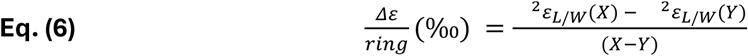

Raw data and code for all data manipulations and statistical analyses is available on GitHub (https://github.com/carolynnharris/Saci_Enviro_d2H) and archived on Zenodo (60)

## 3. Results

We examined how temperature, pH, shaking rate, O_2_ availability, and electron donor (sucrose) flux influence H-isotope fractionation in *S. acidocaldarius* through a series of batch, fed-batch, and chemostat experiments. Details of the growth responses and lipid iGDGT profile responses to these conditions can be found in Cobban et al., (2020) and Zhou et al., (2019). In brief, each environmental condition independently influences growth rate, the relative abundance of many subsets of iGDGT core lipids, and the RI-GDGT (47, 48) (**Table 1**, **FIG 2A-E, S1**).

**Fig. 2.**
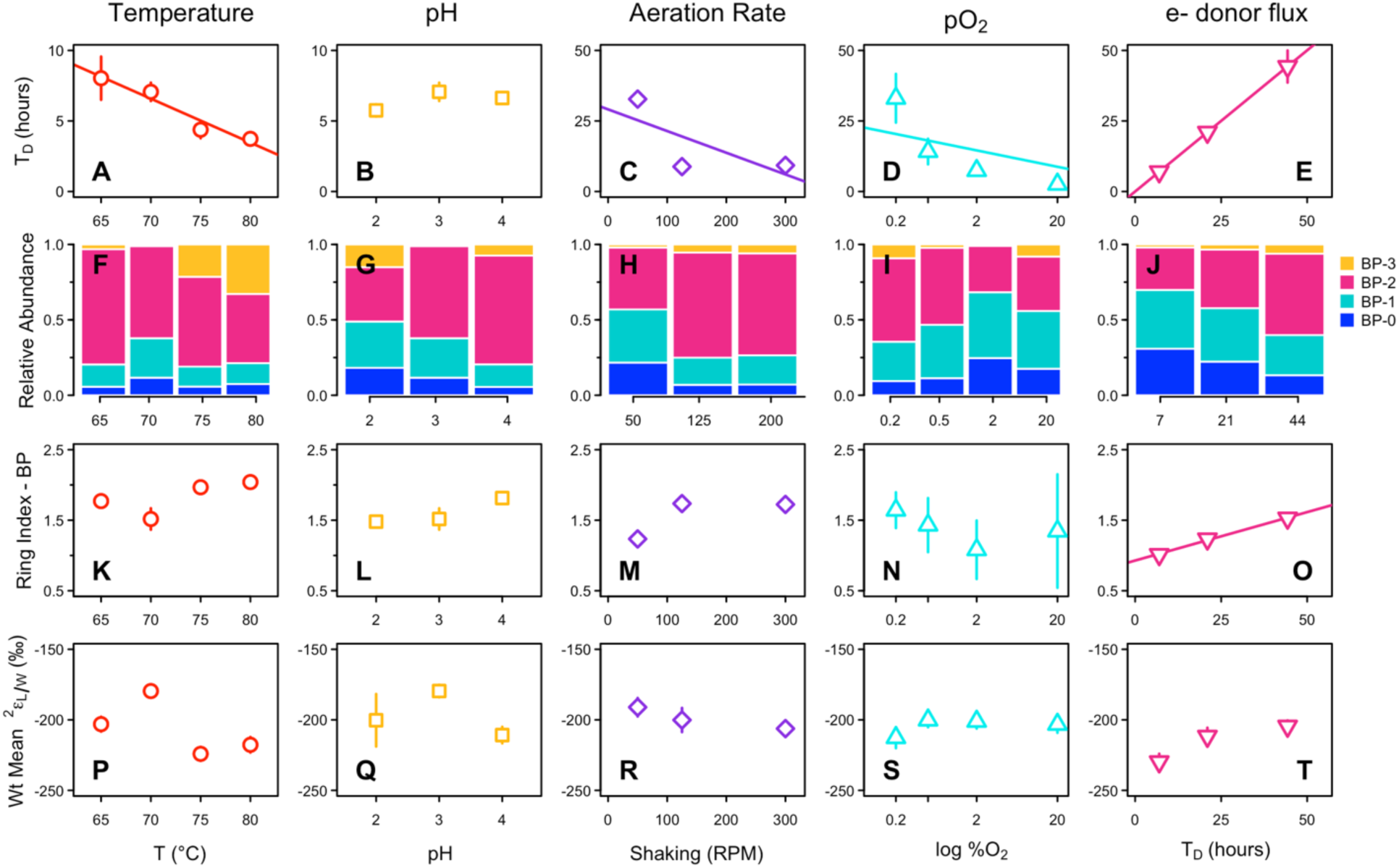
Results for *S. acidocaldarius* grown under varying environmental conditions. Top row (**A-E**): Doubling time in response to each environmental condition. Second row (**F-J**): Relative abundance of biphytane (BP) lipids derived from iGDGTs. BP-3 was the least abundant and was below the detection limit in some samples. Bars represent the mean across replicates for each treatment level. Third row (**K-O**): Ring Index (RI-BP) of biphytane lipids derived from iGDGTs. Bottom row (**P-T**) The abundance weighted mean H-isotope fractionation (^2^ε_L/W_) between growth water and biphytanes. For all rows, points represent the mean and propagated error across biological and technical replicates for each treatment level. In many cases, error bars are smaller than symbols. Only statistically significant regressions are shown (p < 0.05).

Here, we show that each environmental condition also influences the relative abundance of BP lipids and the RI-BP (**Table 2**, **FIG. 2F-O**). Among all samples, the relative abundance of biphytanes averaged 13%, 30%, 49%, 7% for BP-0, -1, -2, and -3, respectively; BP-3 was the least abundant and was not measurable in some samples. RI-BP ranged from 1.01 to 2.04 and, as expected, was positively correlated with RI-GDGT (R^2^ = 0.52, *p* = 0.001, **FIG. S2**). Among all samples, the abundance weighted mean δ^2^H_BP_ values range from -276 to -225 ‰ and were consistently depleted in ^2^H relative to growth water, corresponding to ^2^ε_L/W_ values from -230 to -180 ‰ (**Table 2**, **FIG. 2P-T**).

**Table 2.** Descriptive statistics for individual biphytanyl (BP) lipids derived from iGDGTs for *S. acidocaldarius* grown under varying environmental conditions. Mean and sd include biological replication (N ≥ 1) and technical replication (e.g., multiple injections, n ≥ 3). Abundance-weighted means are shown for δ^2^H_BP_, ^2^ε_L/W_, and Δε/ring values with propagated error.

Despite significant effects on growth rate and lipid cyclization, individual environmental parameters had minimal impact on ^2^ε_L/W_ values, which spanned a range of 44, 31, 15, 21, and 25 ‰ in the temperature, pH, shaking rate, O_2_ availability, and electron donor flux experiments, respectively. We observed a weak linear relationship between environmental conditions and ^2^ε_L/W_ only in the electron donor flux experiment, where larger doubling times are associated with slightly less negative ^2^ε_L/W_ values (smaller offset from water) (R^2^ = 0.89, *p* = 0.22, slope = 0.72 ± 0.25 ‰/hour) (**FIG. 2T**).

In the batch and fed-batch experiments, ^2^ε_L/W_ values for individual BPs showed no consistent pattern with increasing number of cyclic moieties and the ring difference (Δε/ring) varied widely in direction and magnitude among all treatments (-39 to +24 ‰) (**Table 2**, **FIG. 3A-D, S3A-D**). The variability in mean Δε/ring was smaller, though still not of uniform direction, with values ranging from -5.4 to +10.2 ‰ (**Table 2, FIG. S4A-D**). In contrast, ^2^ε_L/W_ values in the chemostat experiment generally increased with the number of cyclic moieties, showing offsets of 3.7, 4.1, and 10.1 ‰ for BP-1, BP-2, and BP-3 relative to BP-0 (**Table 2, FIG. 3E, S3E**). Similarly, the mean ring difference (Δε/ring) was uniformly positive and showed little variation among doubling times, ranging from 4.9 to 9.2 ‰ (**Table 2, FIG. S4E**). On average in continuous culture, each additional ring contributes a 7.4 ± 2.2 ‰ increase to biphytane ^2^ε_L/W_.

**Fig. 3.**
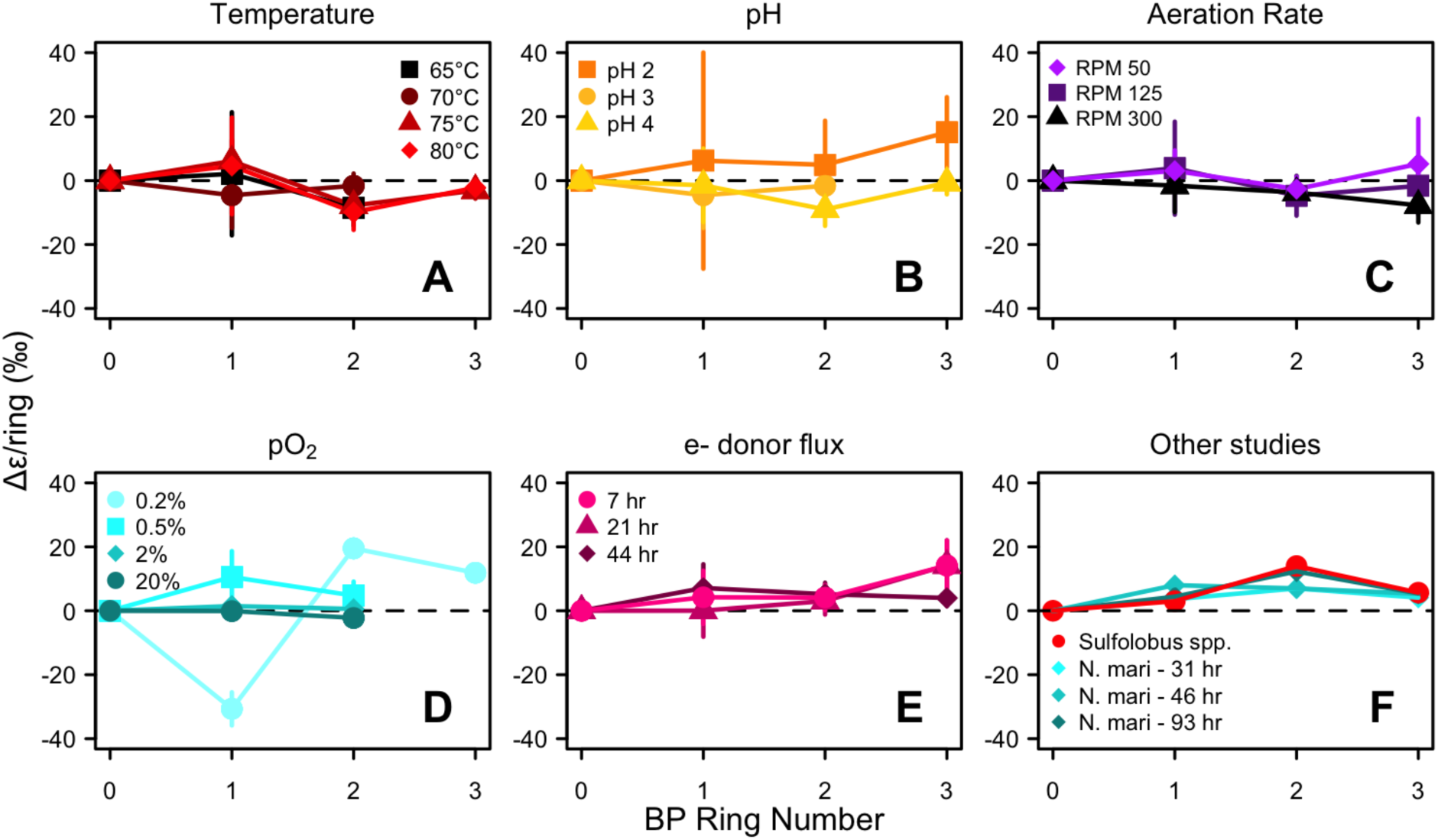
(**A-E**) Ring difference (Δε/ring, ‰) for individual biphytanes in each environmental condition experiment. BP-0 has a Δε/ring of 0 by definition and is shown for reference. BP-3 was not recovered from every treatment. Note the axes are scaled differently for each plot. Points represent mean of biological and technical replicates for each treatment level. (**F**) Ring difference for biphytanes examined in other studies. Blue diamonds are *N. maritimus* from Leavitt et al., 2023; red squares are *Sulfolobus* sp. from Kaneko et al., 2011. Kaneko et al., also recovered BP-4 from *Sulfolobus* spp. (data not shown); the Δε/ring for BP-4 is 11.7 ‰.

Correlation matrices show the linear relationship between each environmental condition, growth parameters (doubling time and maximum OD_600_), mean lipid cyclization indices (RI-BP and RI-GDGT), Δε/ring and abundance weighted ^2^ε_L/W_ values for each environmental condition experiment (**FIG. 4**). These plots further demonstrate that the minimal variation in ^2^ε_L/W_ observed within each experiment is not related to variation in the experimentally manipulated environmental conditions or any other measured parameter.

**Fig. 4.**
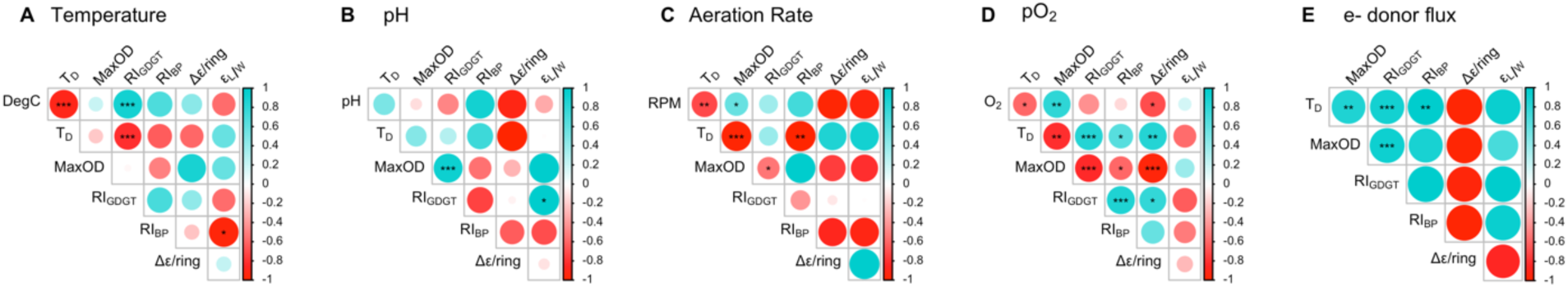
Correlation matrices showing linear relationships among environmentally manipulated parameters, growth statistics (doubling time (T_D_) and maximum OD_600_), mean cyclization of lipids (RI-GDGT and RI-BP), ring difference, and abundance weighted mean ^2^ε_L/W_ for (**A**) temperature, (**B**) pH, (**C**) aeration rate, (**D**) O_2_ mixing ratio, and (**E**) electron donor flux. Notably, mean ^2^ε_L/W_ values do not show a strong linear correlation to any other measured parameter, indicating that environmental conditions, growth statistics, and mean cyclization do not significantly influence the degree of lipid/water fractionation in *S. acidocaldarius*. Circle color and size indicate the strength and direction (blue = correlated, red = anticorrelated) of the relationship. Significance levels: * p < 0.05, ** p < 0.001, *** p< 0.0001. Circles without a star indicate the relationship is not statistically significant.

To explore the possibility that lipid/water fractionation is related to optimal growth, we relate abundance weighted mean ^2^ε_L/W_ values to doubling time for each environmental condition (**FIG. 5A-E**). To allow for comparisons among all environmental conditions, we also relate ^2^ε_L/W_ to doubling time for all experiments combined. In this pooled dataset, abundance weighted mean ^2^ε_L/W_ values range from -230 to -180 ‰ (mean = -204 ± 12 ‰, based on 22 biological replicates; **Table 2**) and showed no correlation to doubling time (**FIG. 5F**, R^2^ = 0.0, *p* = 0.97), indicating a decoupling between lipid/water fractionation and growth rate.

**Fig. 5.**
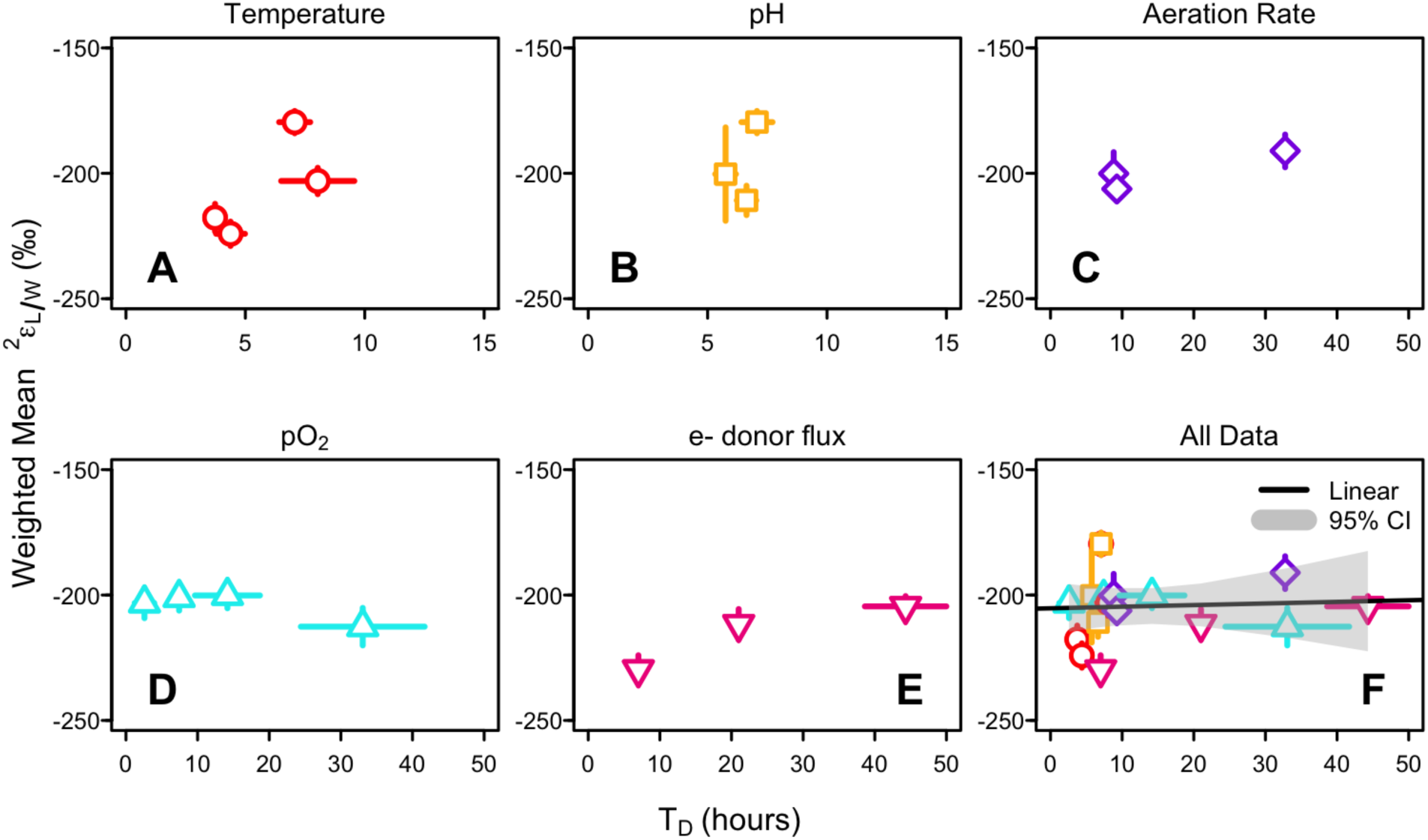
The abundance weighted H-isotope fractionation (^2^ε_L/W_) between growth water and biphytanes (BPs) in response to doubling time (hours) in each individual environmental condition experiment (**A-E**) and pooled across all experiments (**F**). When data from all environmental control experiments is pooled, there is no relationship between doubling time (hours) and abundance weighted mean ^2^ε_L/W_. Points represent mean of biological and technical replicates for each treatment level. Error bars represent propagated error associated with BP abundance measurements, BP-δ^2^H measurements, and weighted mean ^2^ε_L/W_ calculations; bars are smaller than the symbols in many cases.

## 4. Discussion

### 4.1 Effect of environmental conditions on archaeal lipid ^2^ε_L/W_ values

The primary goal of this work was to investigate the impact of environmental conditions on lipid/water fractionation in archaeal lipids, which relates to their potential as archives of hydroclimate across space and time. The conditions tested in this study were chosen to encompass the natural conditions of thermoacidophilic archaea, which are globally distributed in hot, acidic features including fumaroles (steam saturated discharges) as geysers and pools in the terrestrial environment, and hydrothermal vents in the marine environment (61). Both realms experience steep and rapidly changing gradients in temperature, pH, oxygen, and dissolved solute concentrations, which can disturb ecological niches (62). As a result, Archaea have adapted to or evolved strategies to mitigate these challenges (61, 63).

In *S. acidocaldarius*, our experimental manipulations of temperature, pH, shaking rate, electron acceptor (O_2_) availability, and electron donor (sucrose) flux elicited a 20-fold change in growth rate and 2-fold change in mean biphytane cyclization (RI-BP), but ^2^ε_L/W_ values were largely invariant (-204 ± 12 ‰, n = 22). This finding is consistent with previous studies that show archaeal lipids are consistently ^2^H-depleted relative to source water and display a narrow range of ^2^ε_L/W_ values despite considerable variation in taxonomy, metabolic mode, and growth conditions (10–15) (**FIG 6**).

**Fig. 6.**
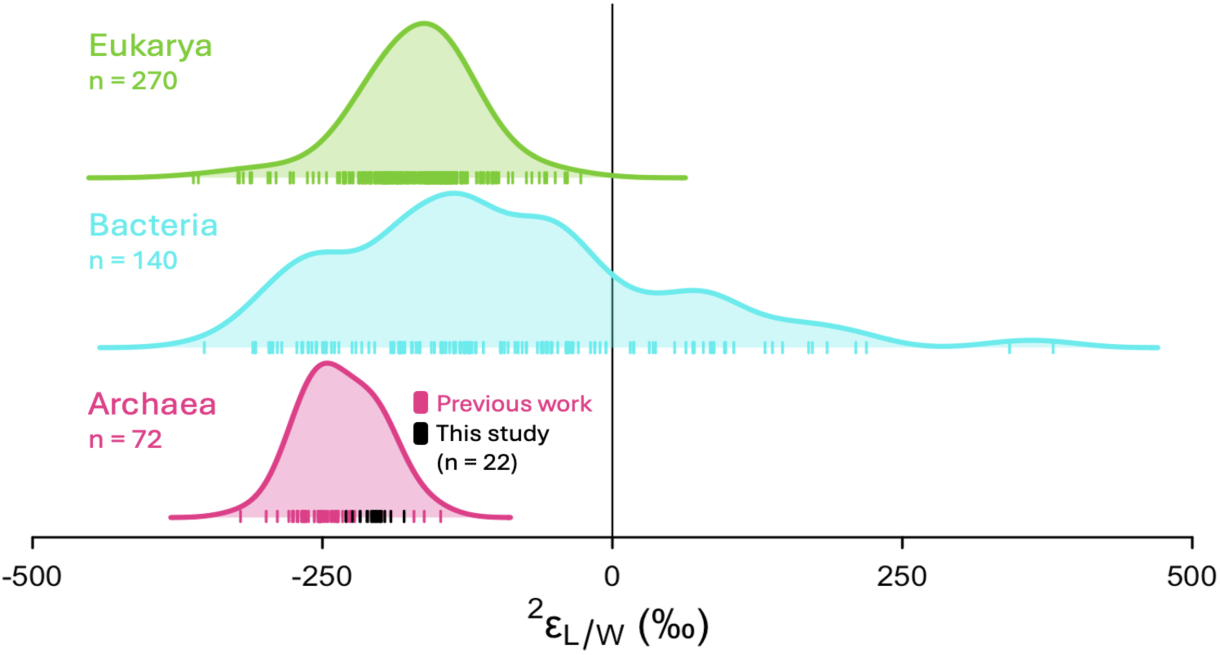
Summary of lipid/water fractionation (^2^ε_L/W_) in culture studies across three domains of life: eukaryotic phototrophs (green; n = 270), bacteria (blue; n = 140), and archaea (pink; n = 72). Tick marks indicate individual ^2^ε_L/W_ values; shaded curves indicate probability density functions. Archaea data includes measurements of phytanes and biphytanes from *Sulfolobus* sp. (Kaneko et al., 2011), *M. barkeri* (Wu et al., 2020), *N. maritimus* (Leavitt et al., 2022), *Acidianus* sp. DS80, *A. fulgidus*, and *Metallosphaera sedula* (Rhim et al., 2024), and *S. acidocaldarius* (this study; black tick marks). Data for Eukarya and Bacteria is redrawn from Rhim et al., (2024), Figure 6. Eukarya data includes measurements for acetogenic and isoprenoid lipids. Bacteria data includes fatty acids synthesized by multiple metabolisms (e.g., aerobic and anaerobic heterotrophy, aerobic and anaerobic chemoautotrophy, and photoautotrophy).

In heterotrophic archaea, lipid-bound H can originate from water or the organic substrate used for growth, with intracellular hydride carriers, such as NADPH, mediating the incorporation of H from both sources into lipids (11, 33, 38). In heterotrophs, the NADPH incorporated into lipid products inherits some H from the available organic substrate(s). While there are several potential mechanisms by which each environmental stressor tested here could influence lipid/water fractionation in archaea, our data suggest these processes either do not occur or are insufficient to cause a measurable effect. Temperature, pH, oxygen levels, and nutrient availability are known to trigger physiological responses in archaea, including changes in growth rates, membrane lipid composition, and enzyme activity (47, 48, 52, 64). Enzyme kinetics, which are influenced by growth temperature and oxidative stress (65), could affect lipid/water fractionation if the enzyme catalyzing hydrogen transfer is associated with a significant isotope effect (33). High temperatures and low pH reduce water exchange across archaeal membranes to prevent osmotic stress (64), which could enrich the intracellular water in ²H and potentially reduce apparent lipid/water fractionation (66). Additionally, changes in electron donor availability can shift the pathway by which NADPH is reduced, which could influence the δ²H composition of NADPH, which ultimately supplies protons to lipid products (67, 68). For example, methanogenic archaea switch between the reductive acetyl-CoA and F₄₂₀-dependent hydrogenase systems depending on if electron donors (H_2_ or formate) are abundant or limiting (67). Despite these possible mechanisms, lipid/water fractionation in *S. acidocaldarius* remains largely invariant in each experiment, indicating that these processes either do not occur or are not significant enough to alter fractionation under the tested conditions. An alternative hypothesis is that *S. acidocaldarius* maintains a constant internal metabolic balance, particularly in NADPH production and utilization, regardless of external conditions.

Physiological responses to stressors impact growth rate and examining ^2^ε_L/W_ values in response to optimal vs. suboptimal growth may provide a more coherent explanation for the non-linear patterns we observe across the environmental conditions tested. The ^2^ε_L/W_ values for *S. acidocaldarius*, however, are not correlated to growth rate in any single environmental condition experiment or when data for all conditions are pooled. This finding suggests that cells maintain stable internal metabolic processes resulting in a consistent H-balance over a range of growth rates and is consistent with the theory that a constant lipid/water fractionation is a feature of an imperturbable central energy metabolism (14, 15). *S. acidocaldarius*, like most archaea, employs energy-efficient metabolic strategies (69, 70), possibly as an adaptation to energy-limiting environments (63, 71). In tightly controlled metabolic pathways, the balance between supply and use of NADPH reducing power appears to remain constant, even under energy-stress conditions. Previous research suggests archaea may exist in a permanent state of NADPH deficiency such that all NADPH is immediately used for biosynthesis with little to no flux from NADPH back to NADH (14). This efficiency in NADPH use may explain why changing growth rates do not impact kinetic isotope effects associated with biosynthesis and hydride exchange. NADPH deficiency (deemed NADPH flux imbalance by Wijker et al., 2019) is associated with energy limitation, and can provide a framework to relate patterns in ^2^ε_L/W_ values to the fluxes of H through dehydrogenase and transhydrogenase enzymes. Indeed, NADPH flux imbalance may explain why all archaea (to-date) and autotrophic and obligately anaerobic bacteria display negative ^2^ε_L/W_ values of similar magnitude (15, 33, 36, 38). This similarity is striking given that fatty acids and isoprenoid GDGTs are formed via different biosynthetic pathways and suggests that the ultimate control on ^2^ε_L/W_ values may be related to free energy yields (15).

Both the O_2_ availability and sucrose flux experiments test the hypothesis that energy limitation – either terminal electron acceptor or donor – can alter ^2^ε_L/W_ values in archaea. These experiments produced the largest range in doubling times (2 to 50 hours and 7 to 44 hours in the O_2_ availability and electron donor experiments, respectively); yet ^2^ε_L/W_ values remained fairly consistent (-217 to - 196 ‰ and -276 to -244 ‰, respectively). *N. maritimus* is the only other archaeon in which the relationship between electron donor (ammonium) flux and lipid/water fractionation was examined; this archaeon similarly exhibited no significant correlation between a three-fold change in doubling time and ^2^ε_L/W_ values (R^2^ = 0.22, *p* = 0.07, slope = 0.18 ± 0.09 ‰/hour) (14). The authors speculate this strain maintains a consistent H-balance over a range of electron donor fluxes as an adaptation that allows cells to perform nitrification with energy-efficient CO_2_ fixation in oligotrophic environments (14). These findings suggest that archaea with diverse physiologies (e.g., heterotrophy vs. chemoautotrophy, extremophily vs. mesophily) maintain a constant lipid-to-water fractionation that is minimally affected by free energy availability and may reflect the fact that Archaea are adapted to chronic energy limitation (63, 71). This stability may reflect tightly regulated physiological processes, such as isoenzyme switching and modulation of enzyme activities, which help maintain metabolic homeostasis under both optimal and sub-optimal growth conditions (72). In contrast, in bacteria and some planktonic eukaryotes faster growth is associated with larger lipid/water fractionation (66, 73, 74) and ^2^ε_L/W_ values for Bacteria are influenced by electron donor flux (33, 36, 38).

The only other study in archaea to examine the relationship between environmental conditions and lipid/water fractionation cultured the halophilic heterotrophic archaeon, *H. marismortui* (which produces archaeol diethers), under varying temperatures and salinities when grown on complex rich medium (11). The resulting ^2^ε_Archaeol/W_ values are relatively invariant (ranging from -126 to -143 ‰; more enriched than our observations for *S. acidocaldarius*) and are decoupled from growth rate (**FIG. S5;** R^2^ = 0.07, *p* = 0.7, slope = 0.85 ± 1.84 ‰ / hour), providing more evidence that archaea maintain a stable internal H-balance across a range of growth conditions (11).

The impact of growth phase on H-isotope dynamics has not been studied in depth. The batch and fed-batch experiment techniques used here for *S. acidocaldarius* and in the *H. marismortui* study integrate across growth stages, whereas the chemostat electron donor flux experiments for *S. acidocaldarius* and *N. maritimus* achieve constant growth rates. A recent study grew *A. fulgidus* in batch cultures and found that ^2^ε_L/W_ values were more depleted (larger offset from water) during exponential phase relative to stationary phase (15). This pattern was consistent between heterotrophic and autotrophic experiments. The culture conditions in Rhim et al. resulted in a threefold difference in growth rates (similar change to the *N. maritimus* study, but much smaller range than the 20-fold change we report for *S. acidocaldarius* here) and a 50 ‰ change in ^2^ε_L/W_ values (-280 ‰ to -230 ‰). Rhim and coauthors conclude that both metabolic mode (heterotrophic vs. autotrophic growth) and growth phase impact ^2^ε_L/W_ values, though the total variance among all treatments was small and overlaps which ranges reported for other archaea (15).

### 4.2 Effect of biphytane ring number on archaeal lipid ^2^ε_L/W_ values

Stoichiometric modeling predicts that BPs with more rings incorporate less of the highly-fractionated hydrogen donated through GGR reduction, a process where the enzyme geranyl-geranyl reductase (GGR) catalyzes the reduction of double bonds in isoprenoid chains, thereby affecting the isotopic composition of the resulting lipids (14). Interestingly, in our *S. acidocaldarius* work, this stepwise enrichment is only seen in the chemostat experiments, where each additional ring contributes 7.4 ± 2.2 ‰ to biphytane ^2^ε_L/W_ values. In the batch and fed-batch experiments, there is no consistent dependence of ^2^ε_L/W_ values on the number of biphytane moieties. In particular, the Δε/ring trends differ between the shaking rate and O_2_ availability experiments, which likely reflects the fact that shaking rate can also impart stress via physical agitation in addition to increasing oxygenation of microbial media (47).

iGDGT-derived BPs extracted from marine sediments also lack a clear stepwise enrichment with increasing ring moieties (10), but this likely occurs because sediments integrate lipids synthesized by different populations of archaea that are either growing on multiple substrates or in different water masses (10). It is less clear why the pure *S. acidocaldarius* batch and fed-batch cultures do not display a ring-specific pattern, though it perhaps relates to variations in biosynthetic fractionations accompanying changes in lipid biosynthesis across the growth phases (44). Considering all the available data, this enrichment pattern appears only minimally affected by differences in growth rate and metabolism, as the magnitude of per-ring increase in ^2^ε_L/W_ value is similar for *Sulfolobus* sp. (10), *S. acidocaldarius* (chemostat experiment in this study), and *N. maritimus* (14), which collectively represent a 5-fold range of growth rates, and a variety of metabolisms and carbon substrates (*Sulfolobus* sp. = yeast extract + O_2_; *S. acidocaldarius* = sucrose + O_2_; *N. maritimus* = chemoautotroph on NH_4_ + O_2_). The magnitude of this stepwise enrichment, however, is small and would likely have a minor impact (usually < 10 ‰) on the abundance weighted mean ^2^ε_L/W_ value even in cases where the BP Ring Index was markedly different among samples.

### 4.3 Archaeal lipid ^2^ε_L/W_ as a function of central metabolism

The *S. acidocaldarius* ^2^ε_L/W_ values we report are broadly similar to results from previous experimental investigations of the *Sulfolobus* genus cultured heterotrophically. When grown at 75 °C and pH 2 on yeast extract, a *Sulfolobus* strain displayed ^2^ε_L/W_ values from -213 to -161 ‰ for BP-0 to -4 (abundance weighted mean = -180 ‰ (10) vs. -204 ± 12 ‰ for *S. acidocaldarius* here). In another study, *Sulfolobus solfataricus* grown at 76 °C and pH 3.5 on glutamate displayed more enriched ^2^ε_L/W_ values, averaging -134 ‰ when intact iGDGT-1 and -2 were analyzed (12). The difference in ^2^ε_L/W_ values may in part be due to the structures analyzed. Lengger and colleagues analyzed intact iGDGTs including the glycerols which may enrich the δ^2^H_iGDGT_ composition, though the different substrate may also play a role (12). Glutamate, an amino acid, is a more complex substrate than simple sugars (such as sucrose) and primarily enters the *Sulfolobus* central metabolism through the TCA cycle (75). Simple sugars enter metabolism primarily through the Entner-Doudoroff pathway, which converts the sugar to intermediates that feed into the TCA cycle (75).

When *S. acidocaldarius* is grown heterotrophically on a simple sugar (glucose), the weighted mean δ^2^H composition of biphytane lipids is strongly correlated with δ^2^H_Water_, indicating a strong first-order control (see *Supplementary Materials*, **FIG. S6**). A simple isotope mass balance model (**Eq. S1**) suggests water contributes at least 56 ± 1% of the total H flux to lipids (**FIG. S7**). Further experimental work and complementary isotope flux models are needed to determine the specific pathways (and associated isotope effects) used during iGDGT synthesis in *S. acidocaldarius* (76).

Autotrophic archaea that produce iGDGTs, however, source their entire H budget to lipids from water in the growth environment and show more negative ^2^ε_L/W_ values than those we observe for *S. acidocaldarius*. In the marine chemoautotroph, *N. maritimus*, grown in continuous culture, BP/water fractionation ranged from -272 to -260 ‰ (14). Similarly, BP/water fractionations for a metabolically-flexible archaeon*, Archaeoglobus fulgidus*, ranged from -280 to -226 ‰ when cultured with several unique carbon substrates and electron donor/acceptor pairs (15). This difference may be driven by the distinct metabolic modes used (e.g., autotrophy vs. heterotrophy) and/or by unique metabolic pathways that employ various intracellular hydride carriers and reductants to bring H’s to the lipid sites, which could generate different net biosynthetic fractionations. Metabolic flux models suggest that in autotrophic archaea (e.g., *N. maritimus*), H-isotope fractionation due to the electron transport chain and NADPH generation result in more depleted lipids than for heterotrophic archaea, such as *Sulfolobus* species, because the source of NADPH impacts ^2^ε_L/W_ values (14). In both heterotrophic and aerobic archaea, most NADPH is generated by glucose dehydrogenase and potentially glyceraldehyde dehydrogenase from the Entner-Doudoroff pathway (72, 77).

Overall, the range of ^2^ε_L/W_ values in Archaeal lipids is narrower than the ranges observed in Bacteria or Eukarya (**FIG. 6**). While the sample size for Bacteria and Eukarya are two-fold and four-fold larger than for Archaea, respectively, Bartlett’s test for homogeneity of variances confirm that Archaeal ^2^ε_L/W_ values have smaller variance compared to Eukarya (K^2^_(1)_ = 16.4, p < 0.0001) and Bacteria (K^2^_(1)_ = 118.5, p < 0.0001) after accounting for sample size. Reported ^2^ε_L/W_ values for bacterial fatty acids vary over 600 ‰ and include large negative and large positive values (15, 33, 36, 38, 78). In Eukarya, ^2^ε_L/W_ values are uniformly negative and span over 300 ‰, though certain lipid classes group together (3, 25, 79). Lipids that are formed via the same biosynthetic pathway (e.g., acetogenic or isoprenoid pathway) tend to occupy distinct ranges of ^2^ε_L/W_ (15). Isoprenoid lipids produced via the MVA pathway in eukaryotes have similar ^2^ε_L/W_ values to iGDGT-derived biphytanes in archaea (15). This similarity suggests that the MVA pathway used for isoprenoid lipid biosynthesis has a large net fractionation in both domains of life.

### 4.4 Applications and Conclusion

Our calibration of biphytane δ^2^H composition as a proxy for the δ^2^H of environmental water in *S. acidocaldarius* has significant implications for paleohydroclimate and paleoecology reconstructions. First, this calibration may enable hydroclimate factors like precipitation, elevation, and seasonality to be reconstructed for environments that do not contain traditionally analyzed lipid biomarkers (e.g., plant waxes) – this includes extreme environments that do not support plant growth (e.g., terrestrial hydrothermal systems, salars, and hyperarid regions). Archaeal lipid δ^2^H values will be most powerful as a proxy in cases where a single-substrate source is a reasonable assumption.

Second, biphytane δ^2^H composition could be applied to marine or lacustrine sedimentary archives where multiple sources of organic matter are suspected, such as a mix of plant waxes representing *in situ* marine/aquatic production and allochthonous terrestrial production (e.g., aeolian deposition). In this case, plant wax δ^2^H values would reflect a mixture of growth waters (marine/lake water and soil water). In contrast, iGDGTs are likely derived solely from archaea living in the overlying water column with few potential terrestrial sources and thus the δ^2^H composition of iGDGTs would only reflect marine/lake water (42).

Here, we show that in the model thermoacidophile *S. acidocaldarius* grown on simple sugars, lipid-water fractionation (^2^ε_L/W_) is not dependent on any of the environmental conditions tested or on growth rate. This finding suggests that *S. acidocaldarius* may have evolved stable metabolisms that exist in a constant NADPH deficit, possibly reflecting adaptation to energy-limiting environments. We conclude that *S. acidocaldarius* will likely demonstrate reliably invariant lipid/water fractionation regardless of growth rate, nutrient status, or energy availability, and that lipid/water fractionation may be more constant for certain archaeons than for phototrophic eukaryotes. For both domains, however, there is more variability in lipid/water fraction relative to the spatial and temporal variability in precipitation δ^2^H values, and more research is needed to understand the sources of this variation so that it can be accounted for in proxy applications.

This calibration underscores the potential for archaeal lipid δ^2^H composition as a proxy for past hydroclimates that could complement or be used independently of eukaryotic lipid δ^2^H composition. Future research should focus on exploring the effects of growth phase and metabolic pathways on lipid/water fractionation in *S. acidocaldarius* and other archaeal species. Expanding these investigations to include more diverse environmental conditions, such as varying carbon substrates or mixed microbial communities, could further validate the consistency of ^2^ε_L/W_ values and refine its utility as a hydrological proxy. Integrating isotope flux models with experimental data such as those generated here, will allow us to better elucidate the specific biochemical pathways responsible for the observed fractionation patterns in domain Archaea.

## Acknowledgements

This research was supported by research grant NSF EAR #1928303940 (WDL, SHK); Simons Foundation Award #623881 (WDL); Dartmouth College (WDL); National Aeronautics and Space Administration (NASA) New Hampshire Space Grant (NASA-SSW-80NSSC19K0539); Deutsche Forschungsge­meinschaft grant 441217575 (F.J.E.). The authors thank the CU Boulder Earth Systems Stable Isotope Lab (CUBES-SIL) Core Facility (RRID:SCR_019300) for analytical contributions; B. Chiu (then at Dartmouth), J. Blewett (then at Harvard), S. Carter (Harvard), and A. Maloney (CU Boulder) for expert laboratory support; Simon D. Kelly from International Atomic Energy Agency, Vienna for measuring the δ^2^H composition of carbon substrates; and all members of the Leavitt lab for all their support.

