## Supplemental Information for "Lipid hydrogen isotope compositions primarily reflect growth water in the model archaeon *Sulfolobus acidocaldarius*"

### Supplementary Information Text

##### S1. Methods for $^2H_2O$-Labeled Water Experiment

To test the incorporation of H derived from water into lipids, we cultured *S. acidocaldarius* on media prepared with either natural abundance water (-49 ‰), with $^2$H-depleted water (-361 ‰; filtered Antarctic ice-core melt water, courtesy of Erich Osterberg) or with $^2$H-enriched water (+415 ‰; achieved via addition of 99.9% $D_2O$) (**Table S1**). Culture media was prepared following the Brock recipe in Wagner et al. (2012), where the complex carbon sources in the media were replaced by 16.7 mmol $L^{-1}$ (0.3% w/v) glucose and 4.0 mmol $L^{-1}$ (0.1% w/v) NZ-amine as the sole organic substrates. Growth was monitored photometrically at regular intervals throughout each experiment by measuring the optical density (i.e., absorbance at 1 cm pathlength) of a culture aliquot at 600 nm ($OD_{600}$) on a spectrophotometer.

Batch experiments were performed in temperature-controlled, shaking incubators (Innova-42, Eppendorf) in 250 mL Erlenmeyer flasks containing 125 mL media covered with loose plastic caps to reduce evaporation. Evaporation was further reduced by maintaining a 2L tray of water in the incubator to humidify the atmosphere. Environmental conditions were 70°C, pH 3, and 100 RPM shaking speed. All experiments were performed in triplicate. Biomass samples were collected in mid-exponential phase by centrifuging (3214 x *g*; Eppendorf 5810 R, S-4-104 rotor) 50 mL of culture for 15 minutes at 4 °C, the resulting cell pellets were stored at -80 °C until lipid extraction. Isoprenoid BP chains were extracted and analyzed as described in the main text. The methods used to determine H-isotope composition of lipids, media waters, and substrates are as described in the main text. $^2$H-labeled waters were diluted with standards of known isotopic composition prior to analysis. The $\delta^2$H composition of glucose was -44.5 ± 2.9 ‰. Based on unpublished data from prior experiments, we expect media water to experience evaporative enrichment of < 3 ‰ over the duration of the experiment.

##### S2. *S. acidocaldarius* H-isotope mixing model

Batch experiments tested the incorporation of water-derived H into biphytane lipids of *S. acidocaldarius*. Doubling times for all treatments ranged from 7 to 11 hours and were unaffected by the $\delta^2$H compositions of growth water (**Table S1, FIG S6A,B**). These rates are consistent with doubling times reported for *S. acidocaldarius* in Cobban et al. (2020) and Zhou et al. (2020). In all samples, BP-1 and BP-2 are the most abundant, and BP-3 is consistently < 10% of the total BP pool; BP distributions were not impacted by $\delta^2H_{Water}$ (**Table S2, FIG S6C**). The $\delta^2$H composition of BPs was strongly correlated to the $\delta^2$H compositions of growth water with a consistent large and negative offset ($R^2 > 0.999$, $p < 0.001$) (**FIG S7A**).

Biphytane $\delta^2H$ values reflect the sum of distinct isotope fractionations between the lipid product and the two potential sources of hydrogen for heterotrophs – water and the organic substrate used for growth. This relationship can be written as an isotopic mass balance equation, originally developed by Zhang et al. (2009), that describes heterotrophic H metabolism as a two-source mixing model (**Eq. S1**). The net fractionations between lipids and each external H source are considered separately in the following model:

**Eq. (S1)**
$$R_L = f_W \cdot {}^2\alpha_{L/W} \cdot R_W + f_S \cdot {}^2\alpha_{L/S} \cdot R_S$$

where $R_L$ is the abundance weighted mean $^2H/^1H$ ratios of BP-0, -1, -2, and -3, $R_W$ is the $^2H/^1H$ ratio of media water, and $R_S$ is the $^2H/^1H$ ratios of the organic substrate (e.g., glucose). $f_W$ and $f_S$ are the fractions of lipid-bound H derived from water and substrate, respectively. $^2\alpha_{L/W}$ represents the net isotope fractionation factor between lipid and water and $^2\alpha_{L/S}$ between lipid and substrate. This model illustrates the overall isotopic relationship between lipids and external hydrogen sources without implying that these sources are directly involved in lipid biosynthesis. For example, glucose may contribute H to lipids through metabolic intermediates, even though it does not directly participate in the biosynthetic reactions.

Our experimental cultures varied $R_W$ while holding $R_S$ constant. The other terms of this model can be constrained by regressing $R_L$ values on $R_W$. The slope of this regression is equivalent to $[f_W \cdot {}^2\alpha_{L/W}]$, and the y-intercept is equivalent to $[(1- f_W) \cdot {}^2\alpha_{L/W} \cdot R_S]$. Because $f_W$ is unknown in heterotrophic growth conditions, we explore the possible range of fractionation factors – $^2\alpha_{L/W}$ and $^2\alpha_{L/S}$ – by varying $f_W$ to range from 0 to 1 (Zhang et al. 2009) (**FIG S7B**). As the fraction of H sourced from water ($f_W$) increases, the expected lipid/water fractionation value ($^2\alpha_{L/W}$) becomes larger and more negative. We can further constrain the likely values of $f_W$ by assuming that lipid/water fractionation is negative ($^2\alpha_{L/W} < 0$; a normal kinetic isotope fractionation). This is a conservative assumption given all previous research on archaea, including the *Sulfolobus* genus, have reported large, negative values for lipid/water fractionation (Kaneko et al. 2011; Rhim et al. 2024). Also, reports for direct water incorporation in bacteria estimate that $^2\alpha_{L/W} = 0.9$ (Zhang et al. 2009; Wijker et al. 2019). With this additional assumption, we can conclude that when *S. acidocaldarius* is grown heterotrophically on a simple sugar, the majority of lipid-H (≥ 56 ± 1 %) is derived from water H, rather than from substrate H.

#### S3. Discussion of H-isotope mixing model

This study aimed to determine the extent to which archaeal biphytane-$\delta^2H$ values record the $\delta^2H$ value of environmental water. In heterotrophic archaea, lipid-bound H can originate from water or the organic substrate used for growth, with intracellular hydride carriers, such as NADPH, mediating the incorporation of H from both sources into lipids (Zhang et al. 2009; Dirghangi and Pagani 2013; Wijker et al. 2019). In heterotrophs, the NADPH incorporated into lipid products inherits some H from the available organic substrate(s). Our experiments show that when *S. acidocaldarius* is grown heterotrophically on a simple sugar (glucose), the weighted mean $\delta^2H$ composition of biphytane lipids is strongly correlated with $\delta^2H_{Water}$, indicating a strong first-order control. Isotope mass balance modeling suggests water contributes at least 56 ± 1% of the total H flux to lipids (**FIG. S7B**). One limitation to this approach is that our experiments were conducted in a semi-defined rich-media, experiments in a defined media using a single carbon source (e.g., glucose only) with $^2H$-labels at particular sites on the molecule will better constrain the relative contribution of substrates and water to lipid products (Harris et al. 2022).

This magnitude of water contribution to lipids in *S. acidocaldarius* aligns with a recent study on the metabolically-flexible anaerobe, *Archaeoglobus fulgidus*, which demonstrates that at least 50% of biphytane-bound H reflects $\delta^2H_{Water}$ when grown heterotrophically, compared to over 80% when grown autotrophically (Rhim et al. 2023). Similarly the aerobic and heterotrophic *H. marismortui* synthesizes diether lipids (archaeol) with $\delta^2H$ compositions tightly correlated with $\delta^2H_{Water}$, indicating a first order control, but $R_{Lipid}$ vs. $R_{Water}$ regression parameters and fractionation constraint curves differ when grown on different carbon sources (Dirghangi and Pagani 2013). Applying the assumption that $^2\alpha_{L/W} < 1$ (a normal isotope effect) suggests that *H. marismortui* derives >78%, >80%, and >12% of archaeol-H from water when grown on pyruvate, succinate, and glucose, respectively. This pattern cannot be explained by a simple inheritance of substrate $\delta^2H$ value (pyruvate = -97‰; succinate = -382‰; and glucose = -61‰) and is likely attributable to large fractionations associated with substrate metabolism and other processes that impact the $\delta^2H$ composition of NADPH (Dirghangi and Pagani 2013).

The large portion of lipid-H that appears to be sourced from water in heterotrophic archaea and in a subset of bacteria (notably heterotrophs in energetically stressful conditions, such as anaerobiosis (Zhang et al. 2009; Dawson et al. 2015; Osburn et al. 2016) could reflect similarities in the physiology and/or ecology of these microbes. Archaea and these bacteria may use common biosynthetic pathways with enzymes that have large isotope effects (Rhim et al. 2024). Overall, this finding is consistent with the narrow range of $^2\varepsilon_{L/W}$ values observed for the Archaea relative to other domains.

### Supplementary Tables

**Table S1**. Summary of culture conditions and descriptive growth statistics for *S. acidocaldarius* grown in media prepared with $^2$H-labeled waters.

**Table S2**. Descriptive statistics for individual biphytanyl (BP) lipids derived from iGDGTs for *S. acidocaldarius* grown in media prepared with $^2$H-labeled waters. Mean and sd include biological replication (N ≥ 1) and technical replication (e.g., multiple injections, n ≥ 3). BP-3 was present in too low abundances to get reliable isotope data. Abundance-weighted means are shown for $\delta^2H_{BP}$ and $^2\varepsilon_{L/W}$ values with propagated error. RI-BP is the BP Ring Index.

### Supplementary Figures

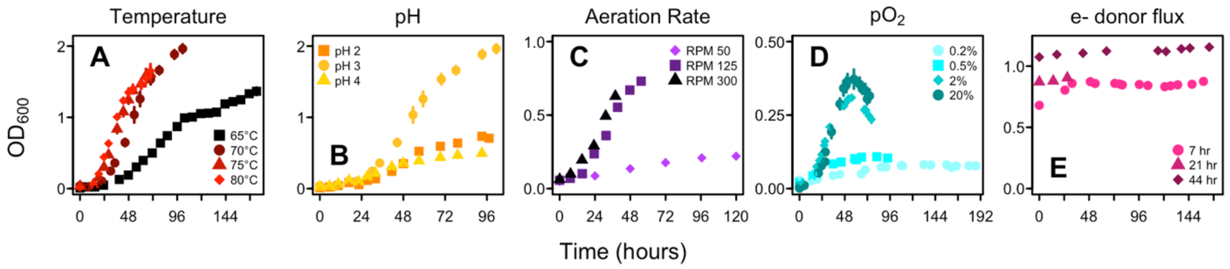


**Fig S1**. Growth curves for *S. acidocaldarius* grown over varying environmental conditions. Originally published in Cobban et al., 2020 (A-D) and Zhou et al., 2020 (E). Points and error bars represent mean and standard deviation of biological replicates for each treatment level. Error bars are smaller than the symbols in many cases.

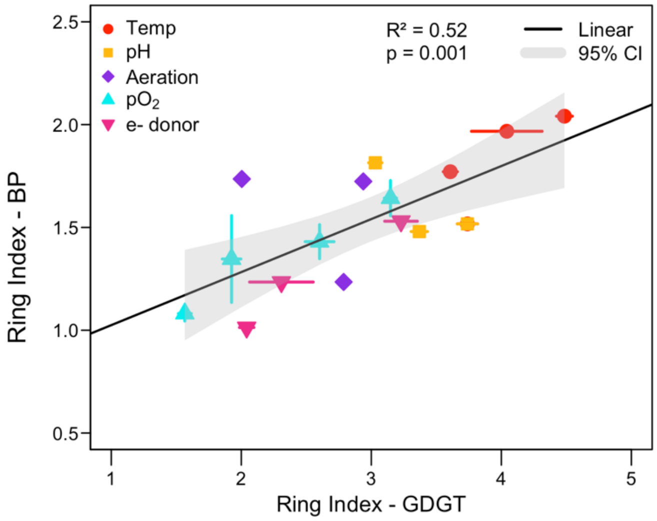


**Fig S2**. Correlation between Ring Indices for parent iGDGTs and derivatized biphytanes across all environmental condition experiments. Points and error bars represent mean and standard deviation of biological and technical replication for each treatment level.

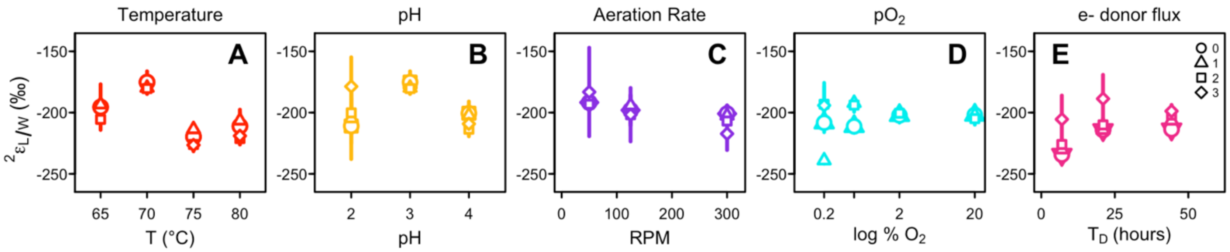


**Fig S3**. $^2\varepsilon_{L/W}$ values (‰) for individual biphytanes (BP-0 = circle; BP-1 = triangle; BP-2 = square; BP-3 = diamond) in response to each environmental condition (A-E). BP-3 was not recovered from every treatment. Points represent means and propagated error across biological and technical replicates.

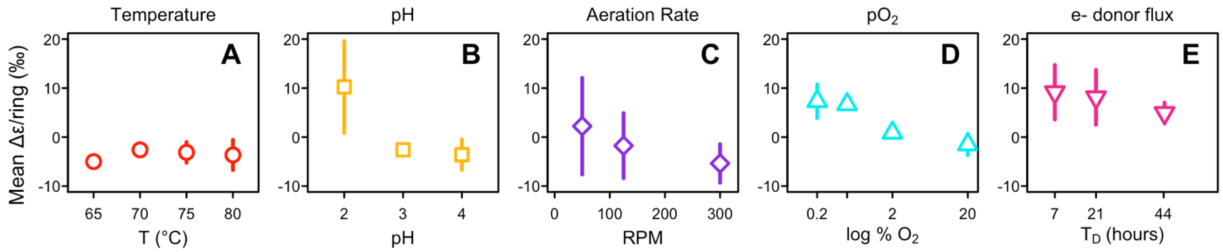


**Fig S4**. Abundance-weighted mean ring difference (Δε/ring, ‰) in response to each environmental condition (A-E). Points show the mean and propagated error across biological and technical replicates.

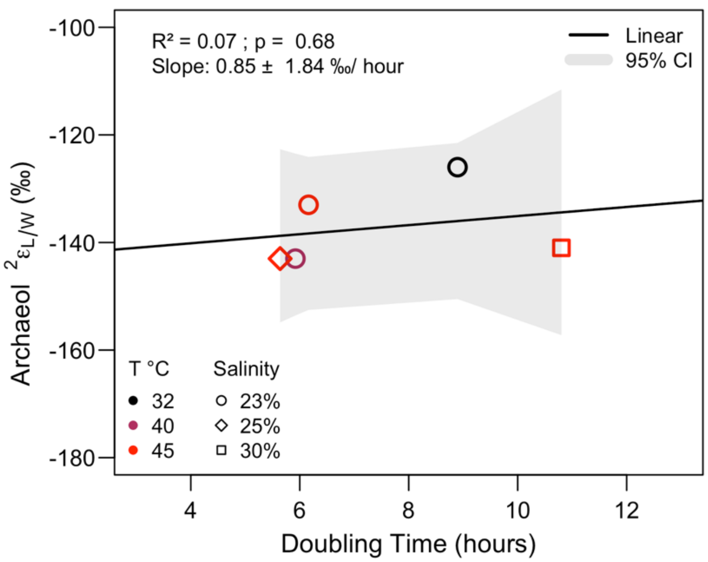


**Fig S5**. The relationship between doubling time and $^{2}\varepsilon_{L/W}$ values for archaeol lipids isolated from experimental cultures of the halophile, *Haloarcula marismortui*, grown over different salinity (symbol shapes) and temperature (symbol color) regimes with yeast extract as the growth substrate (Dirghangi and Pagani, 2013).

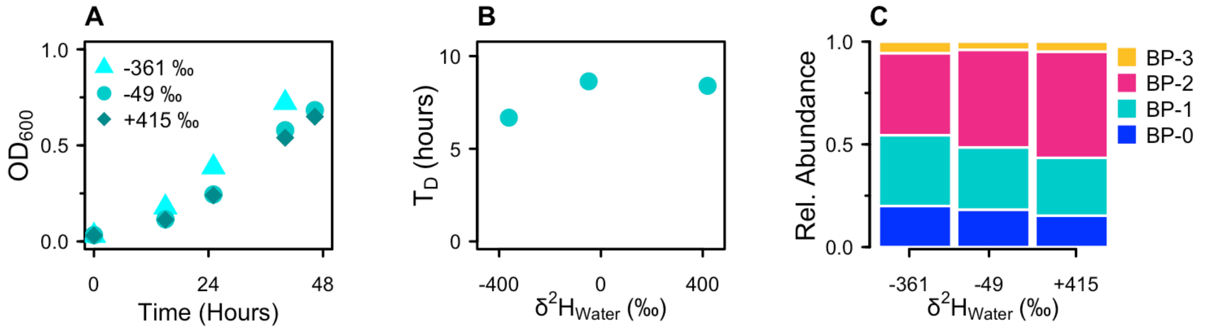


**Fig S6**. A) Growth curves, B) doubling times, and C) relative abundances of biphytane lipids derived from iGDGTs for *S. acidocaldarius* grown in Brock media with varying $\delta^2H_{Water}$ composition. Points and error bars represent the mean and standard error of biological triplicates. Error bars are smaller than the symbols.

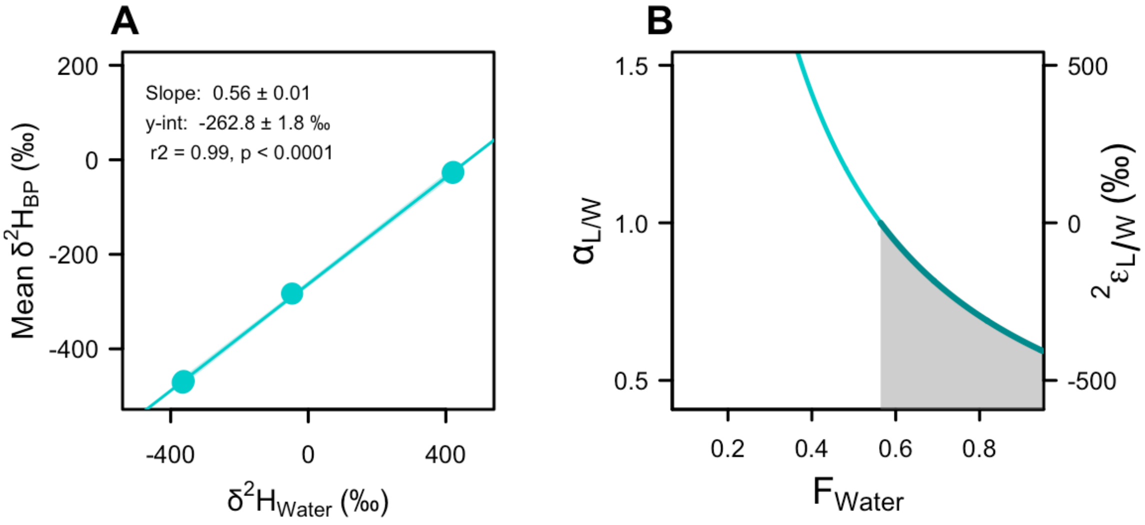


**Fig S7**. Regressions of $R_{Lipid}$ on $R_{Water}$ for the $^2$H-labeled water experiment (A). Points show mean and standard error of biological triplicates. Error bars are smaller than symbols. 95% CI is thinner than the width of the regression line. $R_{Lipid}$ represents abundance weighted mean $R_{BP}$ values. [Note: axes are shown in per mil units for ease of interpretation. Regression parameters were calculated from *R* values]. Because the mass-balance model (Eq. S1) is under constrained, we cannot solve for an exact value of $^2\alpha_{L/W}$; this curve (B) represents the fractionation factor that results from every possible value of $f_{Water}$. 95% CI is thinner than the width of the curve. We can constrain the most likely range of $f_{Water}$ values by assuming a normal kinetic isotope effect where lipids are $^2$H-depleted relative to source water (e.g., $^2\alpha_{L/W} < 1$; shaded region of the curve). To satisfy the assumption that $^2\alpha_{L/W} < 1$, environmental water must supply at least 56 ± 1 % of H flux to lipid products.
