## Supplementary material for "Lipid hydrogen isotope compositions primarily reflect growth water in the model archaeon *Sulfolobus acidocaldarius*": Table 1

**Table 1.** Summary of culture conditions and descriptive growth statistics for S. acidocaldarius grown over varying environmental conditions. $\delta^2H_W$ is the H-isotope composition of media water at the time of lipid sampling. Growth data for batch and fed-batch environmental condition experiments were initially reported in Cobban et al., (2020), growth data for chemostat experiment was initially reported in Zhou et al., (2019). *For batch experiments, RPM is shaking rate; for bioreactor experiments, RPM is impellor speed.

| Experiment | Treatment | N | Culture Conditions | | | | | Growth Rate (hour$^{-1}$) | | Doubling Time (Hours) | | Max $OD_{600}$ | |
|---|---|---|---|---|---|---|---|---|---|---|---|---|---|
| | | | T °C | pH | RPM* | O2 % | $\delta^2H_W$ (‰) | Mean | sd | Mean | sd | Mean | sd |
| Temp (°C) | 65 | 5 | 65 | 3 | 200 | air | -47.6 | 0.09 | 0.02 | 8.03 | 1.53 | 1.37 | 0.12 |
| | 70 | 5 | 70 | 3 | 200 | air | -55.0 | 0.10 | 0.01 | 7.07 | 0.65 | 1.96 | 0.14 |
| | 75 | 5 | 75 | 3 | 200 | air | -54.2 | 0.16 | 0.02 | 4.38 | 0.60 | 1.61 | 0.23 |
| | 80 | 5 | 80 | 3 | 200 | air | -52.7 | 0.19 | 0.01 | 3.73 | 0.19 | 1.65 | 0.21 |
| pH | 2 | 5 | 70 | 2 | 200 | air | -59.8 | 0.12 | 0.01 | 5.75 | 0.43 | 0.71 | 0.04 |
| | 3 | 5 | 70 | 3 | 200 | air | -55.0 | 0.10 | 0.01 | 7.07 | 0.65 | 1.96 | 0.14 |
| | 4 | 5 | 70 | 4 | 200 | air | -60.9 | 0.10 | 0.01 | 6.63 | 0.41 | 0.49 | 0.03 |
| Aeration Rate (RPM) | 50 | 5 | 70 | 3 | 50 | air | -47.6 | 0.02 | 0.00 | 32.75 | 1.50 | 0.22 | 0.01 |
| | 125 | 5 | 70 | 3 | 125 | air | -46.3 | 0.08 | 0.00 | 8.82 | 0.51 | 0.73 | 0.01 |
| | 300 | 5 | 70 | 3 | 300 | air | -46.9 | 0.07 | 0.00 | 9.26 | 0.25 | 0.63 | 0.02 |
| $O_2$ mixing ratio (%) | 0.2% | 3 | 70 | 3 | 200 | 0.2 | -46.9 | 0.02 | 0.00 | 33.05 | 8.64 | 0.08 | 0.00 |
| | 0.5% | 3 | 70 | 3 | 200 | 0.5 | -55.6 | 0.05 | 0.02 | 14.17 | 4.54 | 0.10 | 0.00 |
| | 2% | 3 | 70 | 3 | 200 | 2 | -51.3 | 0.09 | 0.01 | 7.42 | 0.63 | 0.24 | 0.02 |
| | 20% | 3 | 70 | 3 | 200 | 20 | -51.0 | 0.34 | 0.21 | 2.58 | 1.36 | 0.32 | 0.04 |
| e- donor flux ($T_D$, hours) | 7 | 6 | 70 | 2.25 | 200 | 20 | -59.7 | 0.14 | 0.00 | 7.00 | 0.09 | 0.84 | 0.10 |
| | 21 | 9 | 70 | 2.25 | 200 | 20 | -59.5 | 0.05 | 0.00 | 21.00 | 0.45 | 0.88 | 0.03 |
| | 44 | 6 | 70 | 2.25 | 200 | 20 | -50.1 | 0.02 | 0.00 | 44.30 | 5.68 | 1.12 | 0.04 |
