## Supplementary material for "Lipid hydrogen isotope compositions primarily reflect growth water in the model archaeon *Sulfolobus acidocaldarius*": Table 2

**Table 2**. Descriptive statistics for individual biphytanyl (BP) lipids derived from iGDGTs for *S. acidocaldarius* grown under varying environmental conditions. Mean and sd include biological replication ($N \geq 1$) and technical replication (e.g., multiple injections, $n \geq 3$). Abundance-weighted means are shown for $\delta^2H_{BP}$, $^2\varepsilon_{L/W}$, and $\Delta\varepsilon$/ring values with propagated error.

| | | | BP-0 | | | | | | BP-1 | | | | | | | |
|---|---|---|---|---|---|---|---|---|---|---|---|---|---|---|---|---|
| **Experiment** | **Treatment** | **N** | **Rel Abund** | **sd** | **$\delta^2H_{BP}$ (‰)** | **sd** | **$^2\varepsilon_{L/W}$ (‰)** | **sd** | **Rel Abund** | **sd** | **$\delta^2H_{BP}$ (‰)** | **sd** | **$^2\varepsilon_{L/W}$ (‰)** | **sd** | **Δε/ring (‰)** | **sd** |
| Temp (°C) | 65 | 1 | 0.1 | 0.0 | -233.8 | 17.8 | -195.5 | 18.7 | 0.1 | 0.0 | -231.8 | 4.5 | -193.4 | 4.8 | 2.1 | 19.3 |
| | 70 | 1 | 0.1 | 0.0 | -220.6 | 8.4 | -175.2 | 8.9 | 0.3 | 0.0 | -224.9 | 4.9 | -179.7 | 5.1 | -4.6 | 10.3 |
| | 75 | 1 | 0.1 | 0.0 | -262.0 | 4.3 | -219.7 | 4.6 | 0.1 | 0.0 | -256.2 | 4.5 | -213.6 | 4.7 | 6.1 | 6.6 |
| | 80 | 1 | 0.1 | 0.0 | -252.7 | 12.8 | -211.1 | 13.5 | 0.1 | 0.0 | -248.3 | 6.5 | -206.5 | 6.9 | 4.6 | 15.1 |
| pH | 2 | 1 | 0.2 | 0.0 | -258.1 | 25.4 | -211.0 | 27.0 | 0.3 | 0.0 | -252.2 | 19.1 | -204.7 | 20.3 | 6.2 | 33.8 |
| | 3 | 1 | 0.1 | 0.0 | -220.6 | 8.4 | -175.2 | 8.9 | 0.3 | 0.0 | -224.9 | 4.9 | -179.7 | 5.1 | -4.6 | 10.3 |
| | 4 | 1 | 0.1 | 0.0 | -249.2 | 9.0 | -200.6 | 9.6 | 0.1 | 0.0 | -250.6 | 5.9 | -202.0 | 6.3 | -1.5 | 11.5 |
| Aeration Rate (RPM) | 50 | 1 | 0.2 | 0.0 | -230.2 | 3.3 | -191.7 | 3.4 | 0.4 | 0.0 | -227.2 | 5.1 | -188.6 | 5.4 | 3.1 | 6.4 |
| | 125 | 1 | 0.1 | 0.0 | -235.2 | 11.9 | -198.0 | 12.5 | 0.2 | 0.0 | -231.5 | 7.1 | -194.1 | 7.5 | 3.9 | 14.5 |
| | 300 | 1 | 0.1 | 0.0 | -238.4 | 6.5 | -200.9 | 6.8 | 0.2 | 0.0 | -239.9 | 4.6 | -202.4 | 4.8 | -1.6 | 8.3 |
| $O_2$ mixing ratio (%) | 0.2% | 2 | 0.1 | 0.0 | -244.1 | 1.8 | -208.2 | 7.8 | 0.3 | 0.0 | -274.7 | 6.9 | -239.1 | 2.7 | -30.7 | 5.2 |
| | 0.5% | 1 | 0.1 | 0.0 | -254.9 | 5.0 | -211.2 | 5.3 | 0.4 | 0.1 | -237.1 | 4.4 | -192.2 | 4.7 | 10.6 | 8.1 |
| | 2% | 3 | 0.2 | 0.0 | -243.2 | 5.5 | -202.2 | 6.0 | 0.4 | 0.0 | -241.8 | 2.2 | -200.8 | 3.6 | 1.5 | 0.0 |
| | 20% | 3 | 0.2 | 0.0 | -242.6 | 0.5 | -202.0 | 0.7 | 0.4 | 0.0 | -242.7 | 4.2 | -202.0 | 5.0 | -0.1 | 1.5 |
| e- donor flux ($T_D$, hours) | 7 | 1 | 0.3 | 0.0 | -280.0 | 6.7 | -234.3 | 7.1 | 0.4 | 0.0 | -276.0 | 4.0 | -230.1 | 4.3 | 4.2 | 8.3 |
| | 21 | 1 | 0.2 | 0.0 | -261.0 | 5.6 | -214.2 | 5.9 | 0.4 | 0.0 | -260.9 | 5.2 | -214.1 | 5.6 | 0.0 | 8.2 |
| | 44 | 1 | 0.1 | 0.0 | -252.7 | 4.4 | -213.6 | 4.6 | 0.3 | 0.0 | -245.9 | 5.5 | -206.5 | 5.8 | 7.1 | 7.4 |

(Table 2, Part 1) *continues on next page*

| BP-2 | | | | | | | | BP-3 | | | | | | | | Abundance weighted mean | | | | | BP-Ring Index | |
|---|---|---|---|---|---|---|---|---|---|---|---|---|---|---|---|---|---|---|---|---|---|---|
| Rel Abund | sd | $\delta^2H_{BP}$ (‰) | sd | $^2\varepsilon_{L/W}$ (‰) | sd | Δε/ring (‰) | sd | Rel Abund | sd | $\delta^2H_{BP}$ (‰) | sd | $^2\varepsilon_{L/W}$ (‰) | sd | Δε/ring (‰) | sd | $\delta^2H_{BP}$ (‰) | $^2\varepsilon_{L/W}$ (‰) | sd | Δε/ring (‰) | sd | mean | sd |
| 0.8 | 0.0 | -243.3 | 3.0 | -205.5 | 3.2 | -8.6 | 5.5 | 0.0 | 0.0 | - | - | - | - | - | - | -241.0 | -203.1 | 5.3 | -5.0 | 0.0 | 1.8 | 0.0 |
| 0.6 | 0.1 | -225.5 | 3.0 | -180.4 | 3.1 | -1.6 | 3.8 | 0.0 | 0.0 | - | - | - | - | - | - | -224.8 | -179.6 | 4.4 | -2.6 | 0.0 | 1.5 | 0.2 |
| 0.6 | 0.0 | -268.0 | 5.3 | -226.0 | 5.6 | -7.8 | 4.1 | 0.2 | 0.0 | -268.4 | 3.8 | -226.5 | 4.0 | -3.1 | 2.6 | -266.2 | -224.1 | 4.8 | -3.1 | 2.1 | 2.0 | 0.1 |
| 0.5 | 0.0 | -262.2 | 4.6 | -221.2 | 4.9 | -9.9 | 5.5 | 0.3 | 0.0 | -260.1 | 2.7 | -219.0 | 2.9 | -2.2 | 2.7 | -258.9 | -217.7 | 5.5 | -3.6 | 3.1 | 2.0 | 0.0 |
| 0.4 | 0.0 | -248.0 | 10.8 | -200.2 | 11.4 | 4.9 | 13.8 | 0.2 | 0.0 | -227.8 | 22.4 | -178.7 | 23.8 | 15.1 | 11.0 | -248.1 | -200.3 | 18.6 | 10.2 | 9.4 | 1.5 | 0.0 |
| 0.6 | 0.1 | -225.5 | 3.0 | -180.4 | 3.1 | -1.6 | 3.8 | 0.0 | 0.0 | - | - | - | - | - | - | -224.8 | -179.6 | 4.4 | -2.6 | 0.0 | 1.5 | 0.2 |
| 0.7 | 0.0 | -261.5 | 5.5 | -213.6 | 5.8 | -9.0 | 5.1 | 0.1 | 0.0 | -257.3 | 6.5 | -209.2 | 6.9 | -0.7 | 3.6 | -258.9 | -210.8 | 5.9 | -3.6 | 3.1 | 1.8 | 0.0 |
| 0.4 | 0.0 | -231.6 | 4.1 | -193.2 | 4.3 | -2.7 | 3.7 | 0.0 | 0.0 | -222.0 | 34.4 | -183.2 | 36.1 | 5.2 | 14.2 | -229.5 | -191.1 | 6.5 | 2.2 | 9.9 | 1.2 | 0.0 |
| 0.7 | 0.0 | -238.7 | 6.6 | -201.8 | 7.0 | -4.7 | 6.2 | 0.1 | 0.0 | -238.7 | 20.8 | -201.7 | 21.8 | -1.6 | 9.0 | -237.2 | -200.1 | 8.6 | -1.8 | 6.7 | 1.7 | 0.0 |
| 0.7 | 0.0 | -244.1 | 2.1 | -206.9 | 2.2 | -3.7 | 3.2 | 0.1 | 0.0 | -254.0 | 12.7 | -217.3 | 13.3 | -7.7 | 5.4 | -243.5 | -206.2 | 4.4 | -5.4 | 3.9 | 1.7 | 0.0 |
| 0.6 | 0.1 | -231.0 | 13.4 | -193.2 | 17.5 | 19.5 | 2.8 | 0.1 | 0.0 | -230.6 | 9.0 | -194.1 | 15.4 | 11.9 | 1.2 | -248.2 | -212.6 | 7.5 | 7.3 | 3.4 | 1.6 | 0.3 |
| 0.5 | 0.1 | -238.6 | 5.3 | -193.8 | 5.6 | 4.7 | 4.4 | 0.0 | 0.0 | - | - | - | - | - | - | -244.4 | -200.2 | 5.1 | 6.7 | 0.0 | 1.4 | 0.4 |
| 0.3 | 0.0 | -241.5 | 4.1 | -200.5 | 5.0 | 0.6 | 0.2 | 0.0 | 0.0 | - | - | - | - | - | - | -242.0 | -201.0 | 5.2 | 0.9 | 0.9 | 1.1 | 0.4 |
| 0.4 | 0.0 | -245.5 | 4.5 | -205.0 | 5.0 | -2.2 | 0.9 | 0.1 | 0.1 | - | - | - | - | - | - | -243.8 | -203.2 | 6.0 | -1.5 | 2.2 | 1.3 | 0.8 |
| 0.3 | 0.0 | -272.2 | 5.9 | -226.0 | 5.0 | 4.1 | 3.9 | 0.002 | 0.0 | -252.8 | 16.8 | -205.5 | 19.4 | 14.2 | 7.8 | -275.7 | -229.8 | 5.7 | 9.2 | 5.6 | 1.0 | 0.0 |
| 0.4 | 0.0 | -257.1 | 4.7 | -210.1 | 5.0 | 3.1 | 4.2 | 0.003 | 0.0 | -237.0 | 18.4 | -188.7 | 19.5 | 14.2 | 7.9 | -258.6 | -211.7 | 6.1 | 8.1 | 5.6 | 1.2 | 0.0 |
| 0.5 | 0.0 | -241.6 | 2.9 | -202.0 | 3.1 | 5.2 | 3.6 | 0.1 | 0.0 | -238.6 | 4.8 | -198.8 | 5.1 | 4.0 | 2.5 | -244.0 | -204.5 | 4.1 | 4.9 | 2.1 | 1.5 | 0.0 |

(Table 2, Part 2) */end*
