## Supplementary material for "Lipid hydrogen isotope compositions primarily reflect growth water in the model archaeon *Sulfolobus acidocaldarius*": Table S1

**Table S1.** Summary of culture conditions and descriptive growth statistics for *S. acidocaldarius* grown in media prepared with $^2$H-labeled waters. RPM is shaking rate.

| | | | Growth Conditions | | | | Growth Rate (hour$^{-1}$) | | Doubling Time (Hours) | | Max OD | |
|---|---|---|---|---|---|---|---|---|---|---|---|---|
| Experiment | $\delta^2H_W$ (‰) | N | T °C | pH | RPM* | $O_2$ % | Mean | sd | Mean | sd | Mean | sd |
| Water Label | -362 | 3 | 70 | 3 | 200 | air | 0.10 | 0.00 | 6.68 | 0.10 | 0.72 | 0.01 |
| | -48 | 3 | 70 | 3 | 200 | air | 0.08 | 0.00 | 8.64 | 0.27 | 0.68 | 0.01 |
| | +419 | 3 | 70 | 3 | 200 | air | 0.08 | 0.00 | 8.40 | 0.10 | 0.65 | 0.01 |
