## Supplementary material for "Lipid hydrogen isotope compositions primarily reflect growth water in the model archaeon *Sulfolobus acidocaldarius*": Table S2

**Table S2**. Descriptive statistics for individual biphytanyl (BP) lipids derived from iGDGTs for *S. acidocaldarius* grown in media prepared with $^2$H-labeled waters. Mean and sd include biological replication (N ≥ 1) and technical replication (e.g., multiple injections, n ≥ 3). BP-3 was present in too low abundances to get reliable isotope data. Abundance-weighted means are shown for $\delta^2H_{BP}$ and $^2\varepsilon_{L/W}$ values with propagated error. RI-BP is the BP Ring Index.

| | | | BP-0 | | | | | | BP-1 | | | | | | BP-2 | | | | | |
|---|---|---|---|---|---|---|---|---|---|---|---|---|---|---|---|---|---|---|---|---|
| **Experiment** | **$\delta^2H_W$ (‰)** | **N** | **Rel Abund** | **sd** | **$\delta^2H_{BP}$ (‰)** | **sd** | **$^2\varepsilon_{L/W}$ (‰)** | **sd** | **Rel Abund** | **sd** | **$\delta^2H_{BP}$ (‰)** | **sd** | **$^2\varepsilon_{L/W}$ (‰)** | **sd** | **Rel Abund** | **sd** | **$\delta^2H_{BP}$ (‰)** | **sd** | **$^2\varepsilon_{L/W}$ (‰)** | **sd** |
| | -362 | 2 | 0.2 | 0.0 | -469.1 | 2.7 | -165.8 | 1.6 | 0.3 | 0.0 | -467.0 | 0.6 | -162.4 | 1.8 | 0.4 | 0.0 | -472.3 | 3.2 | -170.8 | 2.4 |
| Water Label | -48 | 1 | 0.2 | 0.0 | -288.6 | 5.5 | -253.5 | 5.8 | 0.3 | 0.0 | -296.9 | 18.6 | -261.2 | 19.6 | 0.5 | 0.0 | -298.8 | 13.1 | -263.2 | 13.7 |
| | +419 | 2 | 0.2 | 0.0 | -65.3 | 21.1 | -342.1 | 14.3 | 0.3 | 0.0 | -27.5 | 7.6 | -315.5 | 4.7 | 0.5 | 0.0 | -15.5 | 6.0 | -307.1 | 4.8 |

(Table S2, Part 1) *continues below*

| BP-3 | | Abundance weighted mean | | | | BP-RI | |
|---|---|---|---|---|---|---|---|
| **Rel Abund** | **sd** | **$\delta^2H_{BP}$ (‰)** | **sd** | **$^2\varepsilon_{L/W}$ (‰)** | **sd** | **mean** | **sd** |
| 0.1 | 0.0 | -469.7 | 9.1 | -166.7 | 9.1 | 1.3 | 0.0 |
| 0.04 | 0.0 | -283.0 | 4.1 | -247.6 | 4.1 | 1.4 | 0.0 |
| 0.05 | 0.0 | -26.7 | 59.8 | -315.0 | 59.8 | 1.5 | 0.0 |

(Table S2, Part 2) */end*
